# Multifunctional Systems in Synthetic Biology: Single-Stranded DNA Signaling for Precise Control of Gene Activation

**DOI:** 10.1101/2024.10.20.619289

**Authors:** Brian Ng, Yuying Ding, Matthew Cornall, Ravinash Krishna Kumar, Yujia Qing, Hagan Bayley

**Affiliations:** Chemistry Research Laboratory, Department of Chemistry, University of Oxford, 12 Mansfield Road, Oxford, OX1 3TA; Section of Structural and Synthetic Biology, Department of Infectious Disease, Imperial College London, Sir Alexander Fleming Building, London, SW7 2A7

**Keywords:** DNA signaling, selective gene activation, promoter complementation, logic gates, aqueous-in-oil droplets, droplet-interface bilayers

## Abstract

Controllable gene circuits that respond to defined inputs are essential tools in synthetic biology. By leveraging regulatory mechanisms at either transcriptional or translational levels, synthetic responsive systems have been engineered to recognize diverse signals, such as small molecules (e.g., tetracycline) or physical stimuli (e.g., light). However, these approaches have limitations: small-molecule signals often require high concentrations to be effective, and sophisticated engineering is needed to generate responsive effectors. Here, we establish a simple, versatile gene activation system in which short single-stranded DNAs trigger RNA or protein production by complementing defective single-stranded promoters upstream of target genes. We demonstrate selective gene activation with orthogonal promoters, and logic-gate operations with signal pairs. The signaling system operates in compartmentalized nanoliter droplets scaffolded by bilayers. Signal delivery is controlled by selectively disrupting bilayers or applying transmembrane potential to move signals through protein pores, thereby activating genes within the receiver compartments. This work expands the toolset for engineering multifunctional, responsive materials to meet biotechnological and medical needs, enabling gene activation in response to specific cues.

---

In synthetic biology, signaling refers to the transmission of information between biological components to trigger specific processes, such as gene activation. It is a crucial aspect of synthetic biology, enabling engineered biological systems to communicate and respond to their environment.^1,2^ This is particularly important in synthetic cells, which are intricately designed compartmentalized structures typically enclosed by lipid membranes, that can be capable of gene expression. The distinct intracellular environment of these synthetic cells requires precise signaling mechanisms to ensure accurate and timely gene activation.^3,4^ Advances have been made in engineering synthetic cells to communicate both amongst themselves and with their surrounding environment, where communication predominantly relies on a limited set of small-molecule gene inducers activating their corresponding promoters.^5–13^ However, these promoters often still weakly express genes in the absence of the inducer molecule, and rely on simple diffusion of the inducers across membranes (reviewed^2^).

To address these limitations, recent progress in synthetic biology has explored alternative methods using different signals to control gene expression in engineered biological systems, each with distinct advantages and challenges. Translational toehold switches represent an innovative approach in which mRNA forms a secondary structure that blocks ribosome binding until a trigger RNA exposes the ribosome binding site. However, the short shelf life of input RNA remains a key limitation.^14^ Similarly, gene silencing using siRNA is another common strategy, though again the instability of siRNAs limits their long-term effectiveness.^15^

Light-based systems have shown considerable promise in regulating transcription, but they are still constrained by challenges such as phototoxicity and limited light penetration. For example, photocaged T7 RNA polymerase can be activated by UV light to initiate transcription.^16^ In our previous research, we have demonstrated UV-light-activated gene expression by attaching photocleavable streptavidin proteins to promoters, which blocked transcription initiation until UV exposure.^17–19^ However, while UV-light can be used as a precise trigger, it poses a risk of DNA damage due to its toxicity.^20^ Alternatives to UV light, such as split-T7 RNA polymerase controlled by blue light, offer reduced toxicity but require continuous illumination and specialized equipment, which limits their practical use.^21^ Additionally, while efforts to regulate transcription activators and repressors with visible light show promise, they are hindered by high background activity in the absence of light, reducing their overall effectiveness.^22^

Single-stranded DNA (ssDNA) is an another attractive candidate for gene expression control in synthetic biology, due to the stability of DNA molecules and the high specificity of DNA base pairing.^23^ ssDNA signals have been used to trigger various processes, including strand displacement reactions,^24,25^ isothermal amplification reactions,^26^ dissolution of DNA droplets,^27,28^ and control of liposome fusions.^29–31^ An intriguing, yet underexplored, application of ssDNA involves its use in regulating gene expression, which could then control the production of diverse functional RNA and protein products. This capability could have significant implications in fields such as therapeutics and biotechnology, where precise gene regulation is essential for developing targeted treatments and advanced biotechnological solutions.^32^

One approach to using ssDNA in gene activation involves short single-stranded DNAs complementing defective single-stranded promoters located upstream of target genes, thereby triggering RNA or protein production. While this mechanism was initially observed in fundamental studies on RNA polymerases,^33–35^ further application in synthetic cells requires systematic development. In this work, we introduce strategies for selective gene activation using ssDNA signals to complement single-stranded promoters, referred to here as defective genes. Selective gene activation was achieved by using different promoters or by using oligonucleotide inhibitors to remove undesired signals. Additionally, we demonstrated that DNA signals can be split into shorter fragments to create complex logic gates for gene control. We applied this signaling system in nL-sized aqueous-in-oil droplets where droplets were connected by lipid bilayers and thus signals and defective genes could be separated in different droplets. By breaking the bilayer, such as by lowering the solubility of the lipid through oil-exchange,^36^ the signals diffused across the droplets, thereby activating the defective genes. Furthermore, we showed that even without disrupting the bilayer, it was possible to control DNA signal entry through protein pores embedded within the lipid bilayer, by manipulating the membrane potential, thereby demonstrating a method for selective signal entry into different droplet compartments, enabling gene activation in response to specific environmental triggers.

## RESULTS AND DISCUSSION

### DNA signals complement defective genes for selective transcription activation

We designed defective genes that were single-stranded (∼30 nt) in the promoter and the 5’ end of the transcribed region. The defective genes were made by annealing two ssDNA of different lengths (Figure S1). Transcription activation occurred only when the ssDNA signals complemented the promoter region (Figure 1A). To assess the production of RNA, defective genes were designed to code for RNA aptamers that bind and activate the fluorescence of small-molecule dyes. Transcription activation was studied at different signal and defective gene concentrations (both from 0 to 1000 nM), using defective genes coding for the Broccoli RNA aptamer, which activates fluorescence of the molecule DFHBI-1T (chemical structures and full names are shown in Figure S2).^37^ After 3 h of mixing signals with defective genes, a 35-fold increase in Broccoli fluorescence was observed with as little as 10 nM of a defective gene coding for Broccoli (sequence B1, DNA sequences are in Table S1), and 10 nM single-stranded DNA signal (sequence s1) complementary to the defective Broccoli (Figure 1B, Supplementary Text 1). Furthermore, background fluorescence without the defective gene or the signal remained within 2-fold of the fluorescence of a control *in vitro* transcription (IVT) mixture without any DNA. These results demonstrated the clear responsiveness of our system, with minimal background activity when no signal or defective gene was present in the IVT mixture.

**Figure 1.**
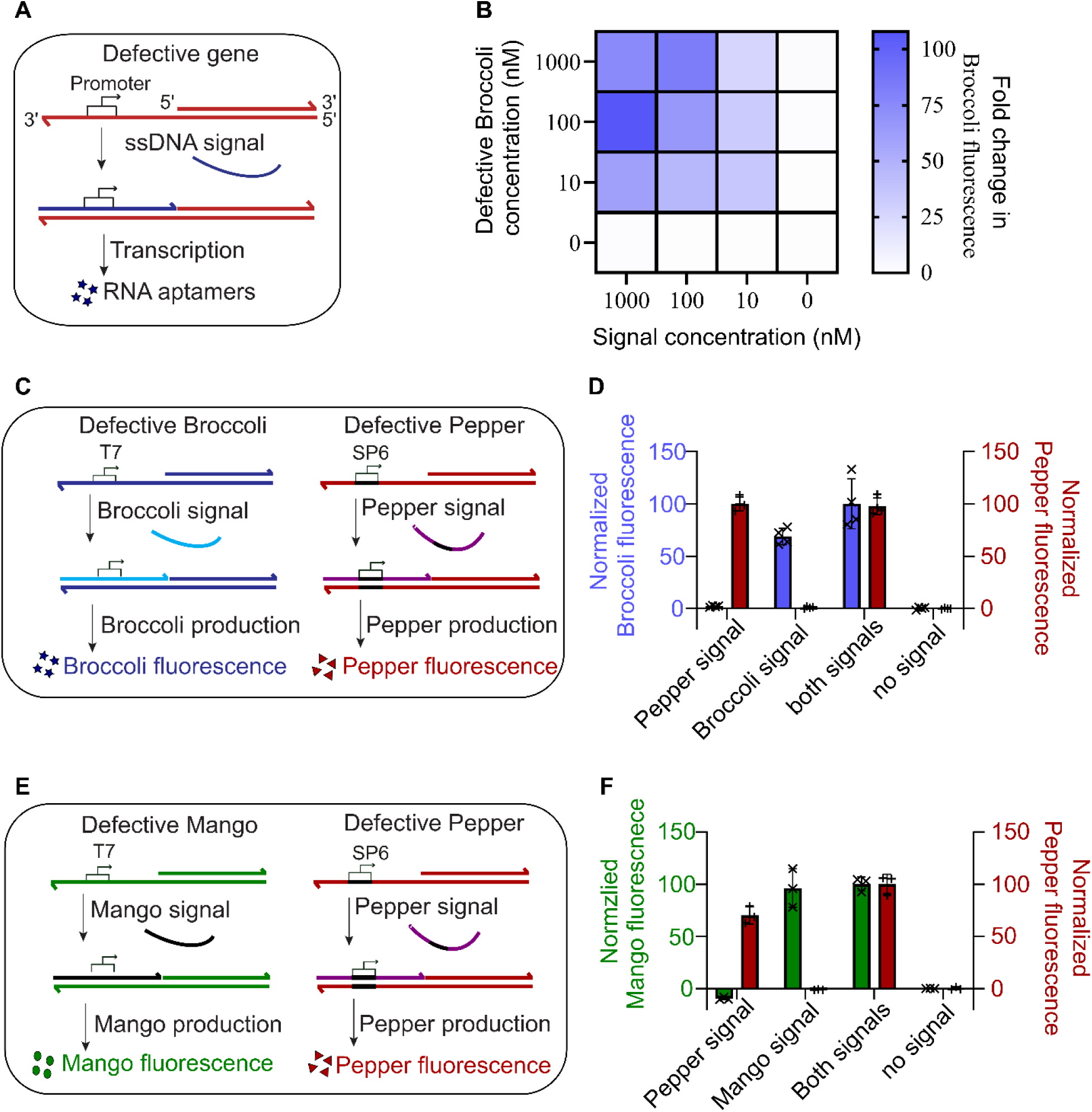
DNA signal complements defective genes for selective transcription activation. (**A**) A schematic of the gene activation process. ssDNA signals complement defective genes with single-stranded promoters to produce RNA transcripts. (**B**) A heat map of fold change in fluorescence intensity from the synthesized RNA in 3 h after different concentrations of defective Broccoli gene (B1) and Broccoli signal (s1) were mixed in an IVT mixture, compared to the fluorescence of the controls with no signal and no defective gene. Values in the heatmap are the mean of n = 4 technical repeats. (**C)** A schematic of selective gene activation using defective Broccoli with T7 promoter and defective Pepper with SP6 promoter. Defective genes are only activated by signals carrying the complementary promoter sequences. (**D**) A bar graph of normalized fluorescence intensity of a defective Pepper (P1, 100 nM) and Broccoli (B1, 100 nM) mixture in an IVT mixture, 3 h after the addition of Pepper (s3, 100 nM) and/or Broccoli (s2, 100 nM) signals. The mean fluorescence intensity of the no-signal control and the brightest sample were normalized to 0 and 100 respectively (detailed methods in supplementary text 2). (**E)** A schematic of selective gene activation using a defective Mango with a T7 promoter and a defective Pepper with an SP6 promoter. Genes are only activated by signals carrying sequences complementary to the defective single-stranded promoters. (**F**) A bar graph of normalized fluorescence intensity of a defective Mango (M1, 100 nM) and Pepper (P1, 100 nM) mixture 3 h after the addition of Pepper (s3, 100 nM) and/or Mango (s4, 100 nM) signals. The mean fluorescence intensity of the no-signal control and the brightest sample were normalized to 0 and 100 respectively. In the bar graphs (**D-F**), technical replicates are displayed by crosses and the heights of the bars are the mean value of the technical replicates. The error bars show the standard deviation.

Similar to small-molecule gene inducers, such as arabinose and IPTG (Isopropyl β- d-1-thiogalactopyranoside) that can activate genes with different promoters,^2^ we sought to determine whether our ssDNA signals could be used for selective gene activation, where different signals would trigger transcription of specific defective genes. To demonstrate this, we engineered genes that encoded different RNA aptamers, which could be distinguished through the binding of different fluorophores (known aptamer-fluorophore pairs reviewed^38^). To achieve this, we investigated the ability of the RNA aptamers Broccoli, Mango and Pepper to activate the fluorescence of DFHBI-1T,^37^ TO1B,^39^ and HBC620^40^ (chemical structures and full names are shown in Figure S2.) in an IVT mixture. We found two sets of orthogonal RNA aptamer-fluorophore pairs: Broccoli/DFHBI-1T and Pepper/HBC620, as well as Mango/TO1B and Pepper/HBC620 (Figure S3, A to C).

Using the orthogonal RNA aptamer system, we developed strategies for selective gene activation using ssDNA signals. In the first strategy, distinct defective genes with different single-stranded promoters were selectively activated by signals carrying complementary promoter sequences. For this, we used an IVT mixture (signal and defective gene concentrations of 100 mM) of defective Broccoli with a single-stranded T7 promoter (B1) and defective Pepper with a single-stranded SP6 promoter (P1) (Figure 1C, S4A). After 3 h, the presence of a Broccoli signal with the complementary T7 promoter sequence (s2) only produced the Broccoli aptamer, while the presence of a Pepper signal with the SP6 promoter sequence (s3) only produced the Pepper aptamer. In the presence of both Broccoli and Pepper signals, both aptamers were produced, whereas without the signals no aptamers were produced (Figure 1D, Supplementary text 2). We also showed the flexibility of this different promoter strategy, which worked for Mango with a defective T7 promoter and Pepper with a defective SP6 promoter. Those constructs had different single-stranded sequences after the promoter, compared to the defective Broccoli (Figure 1, E and F, S4B). Further, we also demonstrated that ssDNA signals can selectively activate genes by using the same RNA polymerase in combination with complementary inhibitors to remove undesired signals when a single promoter is used (Supplementary Text 3, Figure S10-S13).

### Combined use of DNA ligases and RNA polymerases to construct complex logic gates

We next investigated whether by using a combination of DNA ligases and RNA polymerases we could construct complex logic gates from our DNA-activatable transcription system. The Broccoli signal (s2) was split into two shorter ssDNA fragments, each carrying part of the promoter sequence. The downstream piece was 5’-phosphorylated, which allowed the fragments to be ligated by DNA ligases. Transcription activation only occurred when both signal fragments were present, equivalent to an AND gate (Figure 2A). We created four AND gates by splitting the signal after positions −12, −11, −10 and −9 of the transcription start site. Transcription activation was observed in IVT mixtures only when both the upstream signal fragment and the downstream signal fragment were present (Figure 2B). Transcription activation did not occur when the ligase was not present, or when the downstream signal fragment was not 5’-phosphorylated, showing the necessity of ligation for gene activation (Figure 3B, S5).

**Figure 2.**
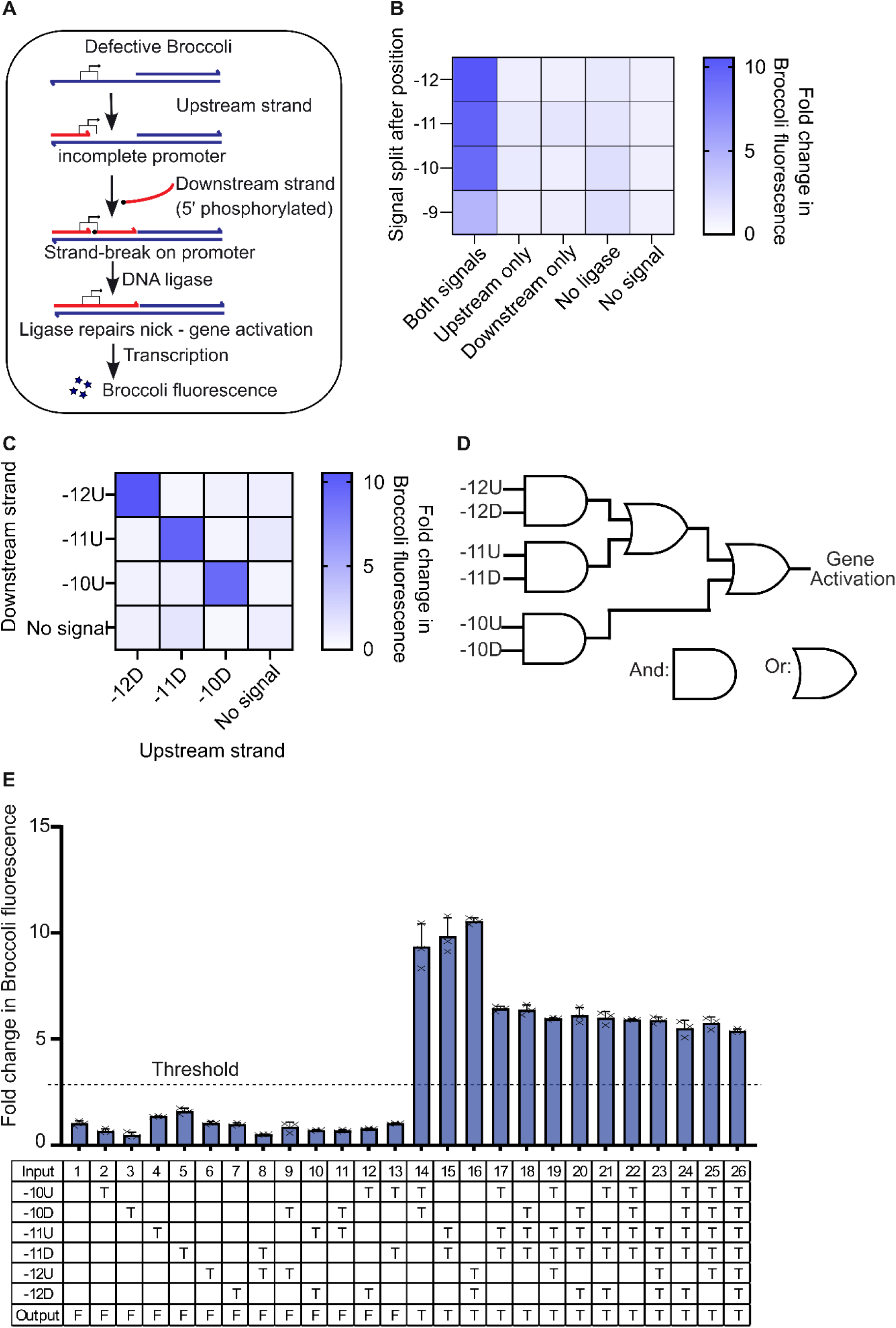
Combined use of DNA ligase and RNA polymerase to effect logic gates. (**A**) Schematic representation of AND gate construction. The DNA signal was split into two shorter ssDNA, each carrying part of the promoter sequence. Transcription activation occurred only when both fragments annealed to the defective gene and the two strands were ligated. (**B)** A heat map of the fold change in fluorescence intensity of the Broccoli aptamer in an IVT mixture 3 h after the defective Broccoli and the signal fragments were mixed. The signal was split after position (B) −12, (C) −11 (D) −10 of the transcription start site, and the outcome compared to the control with no signal fragments. Values in the heatmap are the mean of n = 3 technical repeats. (**C**) A heat map of the fold change in fluorescence intensity of the Broccoli aptamer 3 h after the defective Broccoli gene and signal fragments were mixed in an IVT mixture, compared to the control with no signal fragments. Values in the heatmap are the mean of n = 3 technical repeats. The upstream signal is denoted by U and the downstream signal is denoted by D. (**D**) Equivalent logic gate circuit diagram of an AND-OR-AND-OR-AND gate constructed by combining the AND gates. (**E**) A truth table of AND-OR-AND-OR-AND gates, with a bar graph of the fold change in fluorescence intensity of the Broccoli aptamer in an IVT mixture of the defective Broccoli and different combinations of signal fragments after 3 h, compared to the control with no signal fragments. T = signal fragment that was added. F = signal fragment that was not added. The threshold for activation (dotted line) is set as a 3-fold increase in fluorescence intensity. In the bar graph, technical replicates are displayed by crosses and the heights of the bars are the mean. The error bars show the standard deviation. The signal fragment concentrations were 1000 nM each, and the defective gene concentration was 100 nM.

**Figure 3.**
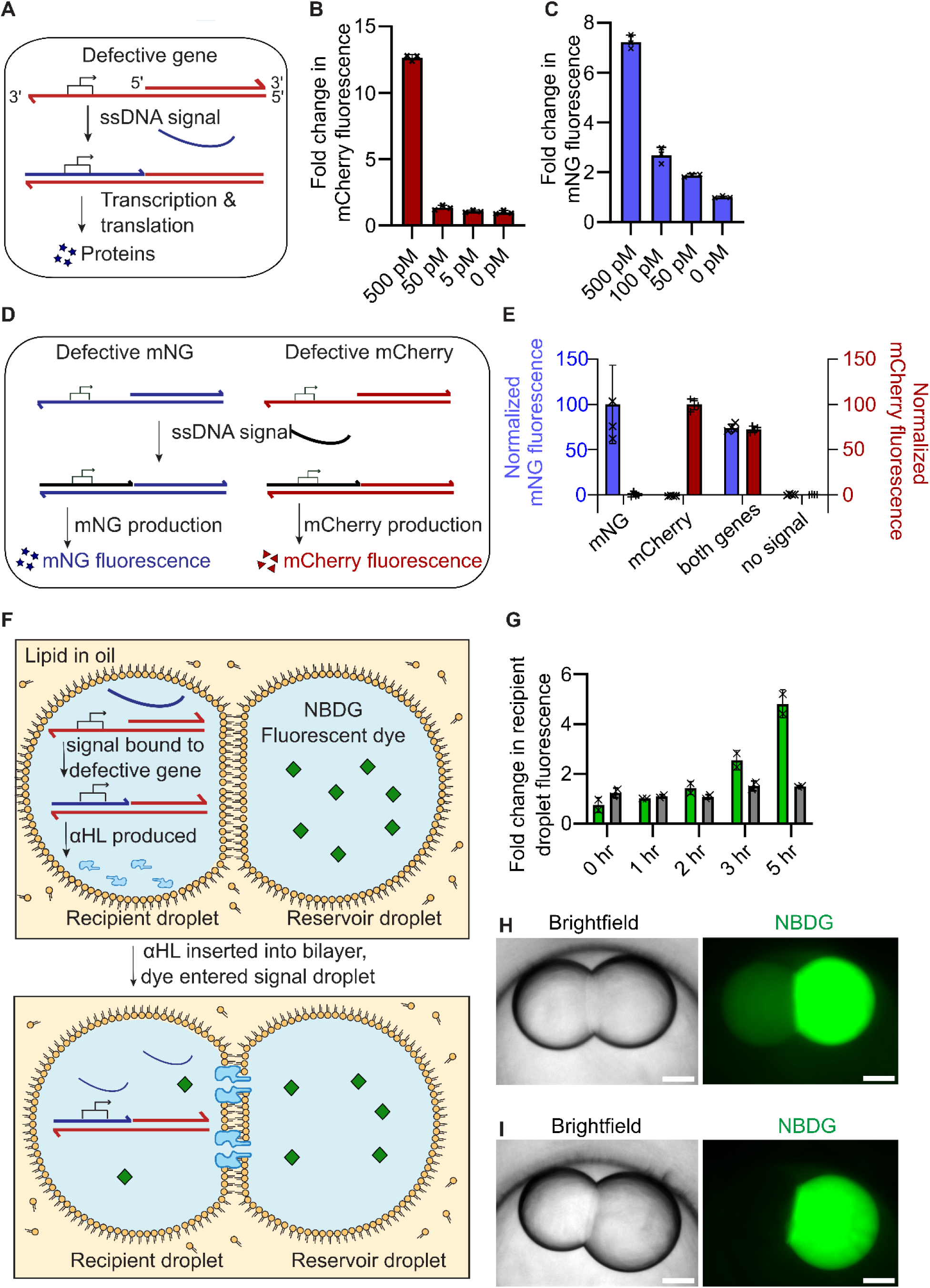
DNA signal triggers functional protein production. (**A**) A schematic of the gene activation process. ssDNA signals complement defective genes with a single-stranded promoter to produce RNA transcripts, which are then translated into proteins. (**B**) A bar graph of the fold change in mCherry fluorescence after 3 h when different concentrations of the DNA signal (s12) were mixed with 5 nM of the defective mCherry gene in an IVTT mixture, compared to the control with no DNA signal. (**C**) A bar graph of the fold change in mNG fluorescence after 3 h when different concentrations of the DNA signal (s12) were mixed with 5 nM of the defective mNG in an IVTT mixture, compared to the control with no DNA signal. (**D**) Schematic of dual protein production by using a universal DNA signal and two different defective genes. (**E**) A bar graph of normalized fluorescence from an IVTT mixture containing the DNA signal (s12) and different defective genes 3 h after signal addition. In each fluorescence channel, the mean fluorescence intensity of the no-signal control and the brightest sample were normalized to 0 and 100 respectively. (**F**) A schematic of a DNA signal activating αHL production in one droplet (left) of a droplet pair. The αHL inserts into the droplet interface bilayer and allows fluorescent dye to enter the recipient left-hand droplet from the right-hand droplet. (**G**) A bar graph of the fold change in fluorescence intensity of recipient droplets overtime, compared to the mean fluorescence intensity at t = 0. Green bar: recipient droplets with the addition of DNA signal (s12); grey bar: recipient droplets without the DNA signal. (**H** to **I**) Brightfield and fluorescence microscopy images of recipient (left-hand) droplets and reservoir (right-hand) droplets after 5 h of incubation. The droplets were in a 1:1 (v/v) ratio of hexadecane to silicone oil AR20, with DPhPC (5 mg mL^-1^) and POPC (5 mg mL^-1^) lipid in the oil. The defective gene concentration was 6 nM, the signal concentration was 20 nM, and the NBDG concentration was 0.33 mM. (**H**) with signal; (**I**) without signal. Scale bars, 200 µm. In the bar graphs (**B**, **C**, **E**, and **F**), technical replicates are displayed by crosses and the heights of the bars are the mean. The error bars show the standard deviation.

We found that the AND gates formed by splitting the signal at −12, −11 and −10 were orthogonal to each other, such that correctly matching pairs of upstream and downstream signal fragments were required for gene activation (Figure 2C). The signal fragments were numbered by the position where the signal was split. The upstream signal fragments were denoted by U and the downstream signal fragments were denoted by D. For example, the signal fragment −10U and the downstream signal −12D contained the entire promoter sequence, however, the pair had overlapping sequences (−12 to −11) and could not be ligated. Subsequently, gene activation was not observed. The three signal pairs in combination created an AND-OR-AND-OR-AND gate, where gene activation was triggered by (−12U and −12D) or (−11U and −11D) or (−10U and - 10D) (Figure 2D). We constructed a truth table of the logic gates using a total of 26 possible combinations of the 6 signal fragments (Figure 2D). When there were more than two fragments present simultaneously, as in this case of −10D, −11U, and −11D (condition 18), the non-matching fragment −10D could compete with the matching fragments −11U and −11D for defective gene binding. Although this competition led to a slight reduction in the final fluorescence intensity (∼20%), the fluorescence of those samples remained clearly distinguishable from IVT reactions lacking any matching pairs of DNA signal fragments (Figure 2E). In these experiments, the signal concentrations were 1000 nM, the defective gene concentrations were 100 nM.

### DNA signal triggers functional protein production

We next investigated whether our ssDNA signals and complementary defective genes could be used to activate protein production in PURExpress *in vitro* transcription and translation (IVTT) systems (Figure 3A). Our designed defective genes coding for proteins such as mCherry and mNeonGreen (mNG) were also made by annealing two ssDNA strands of different lengths. The ssDNA strands were made using a two-step polymerase chain reaction (PCR). A standard PCR was first used to generate a dsDNA template. A linear PCR reaction then made the ssDNA from the dsDNA template, using a single primer and a thermocycling protocol (see Methods). The two ssDNA strands were subsequently annealed to form the defective gene (Figure S6). Protein expression was observed in the IVTT mixture with as little as 500 pM of signal (s12) and 5 nM of a defective gene encoding mCherry (Figure 3B), or 50 pM of signal (s12) and 5 nM of a defective gene encoding mNG (G1) (Figure 3C).

In living cells, transcription factors often activate multiple genes simultaneously.^41^ We next sought to mimic transcription factors using our constructed defective gene system. For example, our DNA signals could mimic transcription factors by simultaneously activating multiple defective genes sharing the same defective promoter. To demonstrate this, we designed a DNA signal (s12) that could complement both defective mCherry (C1) and defective mNG (G1) (Figure 3D). The DNA signal (s12) activated production of mCherry protein in the IVTT mixture in presence of the defective mCherry, activated production of mNG in the presence of the defective mNG, and activated production of both mCherry and mNG in the presence of both defective genes (Figure 3E, S7). In these experiments, the DNA signal concentration was 100 nM, and the total defective gene concentration was 10 nM.

### DNA signal mediates communication in nL-sized aqueous droplets

Finally, we demonstrated the use of our ssDNA signaling system in nL-sized aqueous-in-oil droplet pairs that were connected by a single droplet interface bilayer. The oil composition was a 1:1 (v/v) ratio of hexadecane : silicone oil AR20 that contained 1:1 (w/w) ratio of 1,2-diphytanoyl-sn-glycero-3-phosphocholine (DPhPC) : 1-palmitoyl-2-oleoyl-glycero-3-phosphocholine (POPC) lipid.^42^ We designed defective genes coding for the protein pore α-hemolysin (αHL), a bacterial toxin that lyses cells by forming pores in the plasma membrane.^43^ The defective αHL gene (A1) and the fluorescent dye 2-deoxy-2-[(7-nitro-2,1,3-benzoxadiazol-4-yl)amino]-D-glucose (NBDG) were placed separately in two droplets connected by a droplet interface bilayer (Figure 3F). The lipid bilayer between the droplets is normally impermeable to NBDG. However, in the presence of an activating DNA signal (s12, 20 nM) in the droplet containing the defective gene (6 nM) and IVTT mixture, αHL was produced overtime and inserted into the lipid bilayer, subsequently allowing NBDG (0.33 mM) to diffuse through the protein pores in the membrane. Dye transfer across the lipid membrane was not observed without the DNA signal (Figure 3, G to I).

Next, we explored the physical separation of ssDNA signals from their target defective genes using lipid bilayers. In this setup, the ssDNA signals could only access the defective genes when a suitable environmental stimulus is present. For example, we prepared signal and receiver IVT mixture droplets both formed from a hydrogel (1.5% (w/v) ultra-low-gelling-temperature agarose in a 1:1 (v/v) ratio of hexadecane to silicone oil AR20). The signal droplet contained DNA signal (s4, 70 µM) and the receiver droplet contained the defective Mango gene (M1, 100 nM). By breaking the bilayer, achieved by lowering the lipid solubility through oil exchange to an increased volume fraction of silicone oil,^36^ the signal could diffuse into the receiver compartment and activate Mango aptamer production (Figure 4A). Gene activation only occurred in response to bilayer distruption (Figure 4, B to D).

**Figure 4.**
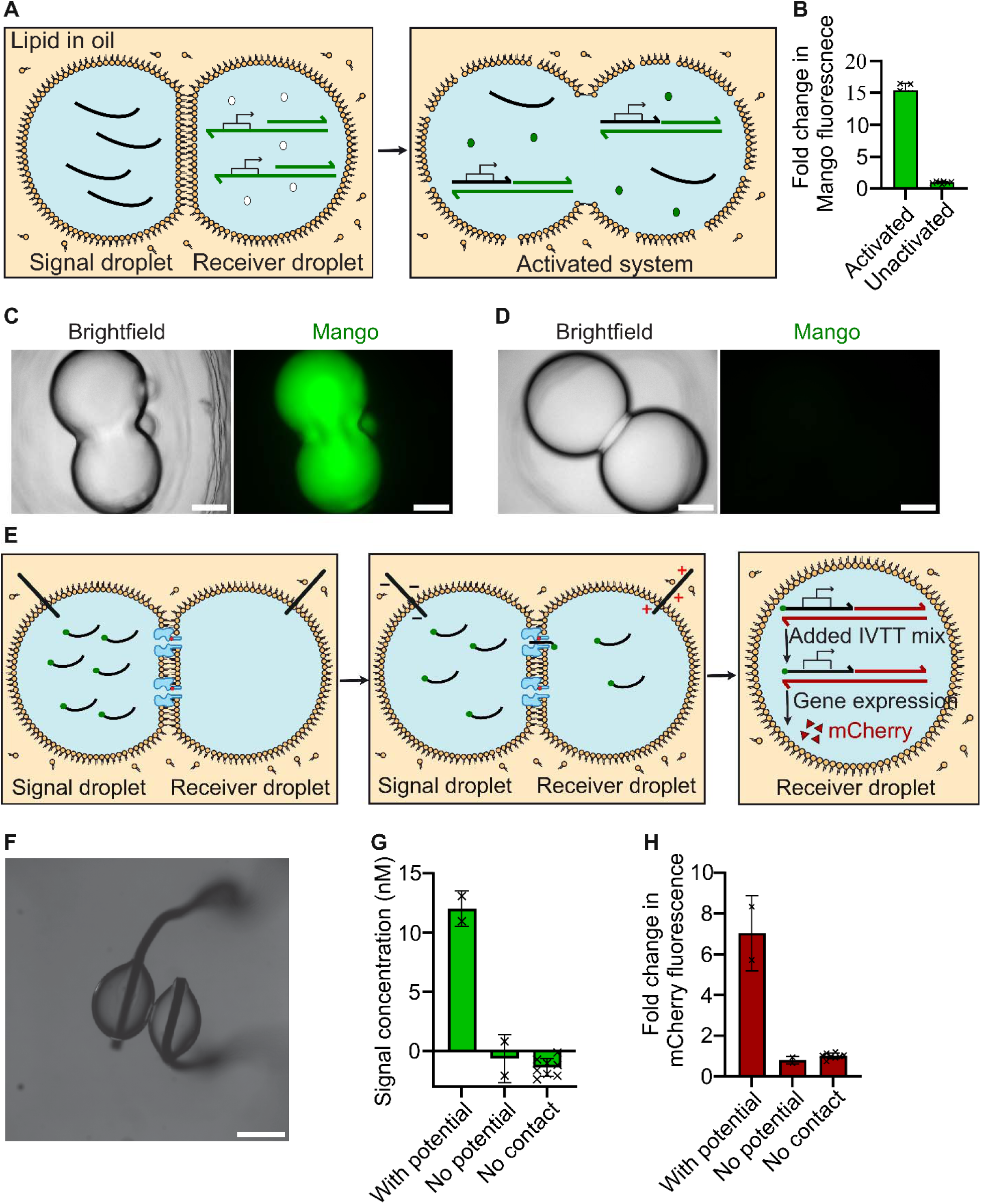
DNA signal-mediated communication in droplets. (**A**) Schematic of gene activation in hydrogel droplets. A lipid bilayer physically separated the signal from the gene expression machinery. Altering the oil to reduce lipid solubility disrupts the bilayer, thereby triggering gene expression. (**B**) Bar graph of the fold change in fluorescence intensity of the Mango aptamer in the hydrogel droplets (1.5% (w/v) ultra-low gelling temperature agarose in a 1:1 (v/v) ratio of hexadecane to silicone oil AR20) after 3 h, compared to the unactivated control. The defective gene concentration was 100 nM, and the signal concentration was 70 µM. (**C, D**) Brightfield and fluorescence microscopy images of hydrogel droplets: one droplet contained the DNA signal (s4) and the other the defective Mango gene (M1). (**C**) Images of an activated system after 3 h. (**D**) Images of an unactivated system after 3 h. (**E**) A schematic of membrane potential controlling DNA signal entry across a bilayer containing synthetic pores. (**F**) Images of a signal droplet and a receiver droplet separated by a droplet interface bilayer. Electrodes were inserted into the droplets to provide the membrane potential. The droplets were in a 1:1 (v/v) ratio of hexadecane : silicone oil AR20, with 10 mg mL^-1^ DPhPC lipid in the oil. The signal droplet contained 300 µM of fluorescently labelled DNA signal (s13) and 0.2 mg mL^-1^ of αHL monomers. Scale bar, 200 µm. (**G**) Bar graph of signal concentrations in receiver droplets with or without an applied potential between the signal droplet and the receiver droplet. (**H**) Bar graph of the fold change in fluorescence intensity of mCherry in receiver droplets after supplementing with IVTT mixture with 5 nM of the defective mCherry gene for 3 h, compared to the control, which has no signal. In the bar graphs (**B**, **G**, and **H**), technical replicates are displayed by crosses and the heights of the bars are the mean. The error bars show the standard deviation.

Next, we explored the use of membrane potential to control DNA signal entry through droplet interface bilayers without disrupting the bilayer. Two aqueous droplets were physically separated by a lipid bilayer (Figure 4, E and F); the signal droplet contained 300 µM of fluorescently-labelled DNA signal (s13) and 0.2 mg mL^-1^ of αHL monomers, which formed pores on the bilayer. The droplets were in a 1:1 (v/v) ratio of hexadecane : silicone oil AR20, with 10 mg mL^-1^ DPhPC lipid in the oil. Although the narrowest diameter of a αHL pore (1.4 nm) is wider than that of ssDNA signal, DNA signal cannot spontaneously translocate through the αHL pores, as the translocation process is entropically unfavourable.^43,44^ We therefore generated a membrane potential through Ag/AgCl electrodes inserted into the droplets. After 1.5 h at 120 mV, 10 nM of signal DNA was transported across the bilayer, as indicated by the fluorescence observed in the droplet from the fluorescently labelled signal (Figure S8). Signal entry was not observed in the absence of a membrane potential over the same time period (Fig 4G). After signal entry into the receiver droplet, the receiver droplet was supplemented with 5 nM of the defective mCherry gene (see methods), along with IVTT components required for protein production. The receiver droplets produced mCherry protein, which led to a 7-times increase in fluorescence intensity compared to the no signal control, while mCherry protein production was not observed in the receiver droplets without the membrane potential (Fig 4H, supplementary text 4).

## CONCLUSION

Our work demonstrates the efficient use of ssDNA signals for gene activation by complementing genes with a single-stranded promoters. Key distinctions of our approach include the selectivity, orthogonality, and modular design of the system, as well as proof of concept for its use in aqueous droplets. By using different promoters or by using oligonucleotide inhibitors to remove undesired signals, we achieved orthogonal gene activation, ensuring that specific signals only activated their corresponding genes without cross-reactivity. The ssDNA signalling triggered the production of multiple RNA transcripts (including aptamers) and proteins. Additionally, we demonstrated that the DNA signals can be split into shorter fragments to construct complex logic gates that control gene activation. The modular nature of our system allows for easy adaptation, enabling the interchange of various promoters and signals to regulate different gene circuits. We applied this signaling system in aqueous-in-oil droplets, where the signals and defective genes were separated by a lipid bilayer. By breaking the bilayer, such as by reducing the lipid solubility through oil exchange, the signal could diffuse across the droplets, thereby activating the defective genes. Furthermore, we showed that even without disrupting the bilayer, it was possible to control DNA signal entry through droplet interface bilayers by controlling the membrane potential, thereby demonstrating a method for selective signal entry into different droplets.

Previous work has made limited use of oligonucleotide complementation to activate genes in synthetic cells.^9,45^ For example, DNA signals were used to activate genes through non-selective diffusion of the DNA signals into polymersomes containing the defective single-stranded promoters,^45^ or by co-encapsulating the DNA signals with the single-stranded promoters in the same synthetic cell.^9^ Our work differentiates itself by addressing selective gene activation using DNA signals, as well as the control of DNA signal movement across lipid bilayers for gene activation, both of which remained unexplored in the earlier studies.

The use of ssDNA as a signaling molecule offers several practical advantages over other methods. DNA sequences are highly programmable, and we showed that as few as 50 pM ssDNA signal was sufficient to activate gene expression, in comparison to small-molecules activators such as IPTG and arabinose where typically µM to mM range of signal concentration is required.^2^ In addition, ssDNA has a low cost and long shelf life.^46,47^ Collectively, these features position our system as an efficient alternative to previous gene activation strategies, particularly in terms of control and applicability in synthetic biology systems.

Our study also expanded our understanding of T7 and related polymerases, particularly their potential in synthetic biology applications. For example, by investigating the effects of splitting ssDNA signals at various positions to create logic gates, we discovered that the T7 promoter is intolerant of strand breaks on the coding strand between positions −12 and −9 relative to the transcription start site. Using this insight, we successfully developed three orthogonal AND gates. Although we did not test whether strand breaks are tolerated in other regions of the promoter, it is plausible that additional orthogonal AND gates could be engineered by introducing strand breaks at other positions.

We showed that the DNA signal and the defective genes could be separated by lipid bilayers in different droplets, and that the defective genes were activated by disrupting the bilayer. While we chose to disrupt the bilayer by changing the oil, lipid bilayers could be made sensitive to other environmental stimuli, including temperature, light, redox states, and enzymes (reviewed^4^). In scenarios where preservation of membrane integrity is desired, we also demonstrated the use of membrane potential to deliver ssDNA signal across protein pores embedded in the bilayer. The membrane potential in this work was maintained by electrodes inserted into the droplets, but it could also be generated by light-responsive proton pumps, asymmetric lipids, or by living cells.^48–50^ The entry of DNA signals into aqueous droplets could be more precisely regulated through site-selective covalent chemistry, ensuring that only signals capable of reacting with the pores can enter specific aqueous droplets.^51,52^ Further, aqueous droplets could be transferred from lipid-in-oil environments to aqueous environments, which would enable direct chemical communication (through DNA signaling) between droplets and their aqueous surrounding.^53^ Looking forward, our work has laid the groundwork for engineering multifunctional droplet networks, where different sets of genes could be specifically activated in response to different DNA signals. This capability could have significant implications in fields such as therapeutics and biotechnology, where precise gene regulation is essential for developing targeted treatments and advanced biotechnological solutions.

## MATERIALS AND METHODS

### Reagents

All reagents were obtained from Merck unless otherwise specified. mCherry, mNeonGreen and αHL plasmids were gifts from the Michael Booth group. T7 RNA polymerase, SP6 RNA polymerase, T3 DNA ligase, T4 DNA ligase, Monarch DNA Gel extraction kit, Monarch PCR cleanup kit, PURExpress, murine RNase inhibitor were purchased from New England Biolabs. DreamTaq Green PCR Master mix and DreamTaq PCR Master mix were purchased from ThermoFisher. All lipids were purchased from Avanti Polar lipids. DFHBI-1T was purchased from Tocris. HBC620 was purchased from MedChemExpress. TO1B was purchased from Applied Biological Materials.

### Defective gene synthesis

Defective genes were made by annealing two ssDNA of different lengths. For short defective genes (<200 nt), DNA strands were purchased commercially. Equal amount of each ssDNA in annealing buffer (100 mM NaCl, 10 mM Tris pH 8.0, 1 mM EDTA) were held at 98 °C for 10 s and cooled at a rate of 0.1 °C s^-1^ to 20 °C. (Veriti 96 Well Thermal Cycler, Applied biosystems).

For longer defective genes (>200 nt), the coding strands were made from a two-step PCR. In the first step, a standard exponential PCR (peqSTAR 2X PCR thermocycler, VWR) was used to make dsDNA template containing the coding strand and it’s complementary strand. The PCR used DreamTaq Green PCR master mix with primers R1 and R2, following the manufacturer’s recommendations. The dsDNA product was purified by agarose gel electrophoresis with Monarch Gel Extraction kit and further purified by Monarch PCR cleanup kit, eluting in nuclease free water. In the second step, linear PCR was used to amplify the coding strand from the dsDNA template. The linear PCR used DreamTaq PCR master mix with primer R4. Unless otherwise specified, [Primer] = 2500 nM, [dsDNA template] = 8 ng µL^-1^, and no. of thermocycles = 20. The ssDNA product was immediately separated from dsDNA template by agarose gel electrophoresis. ssDNA bands were crushed with polypropylene pestle (Bellco glass) in a solution with 300 mM sodium acetate (pH 5.2) and 1 mM EDTA. The mixture was incubated at 37 °C for 16 h, and the gel fragments were removed from the solution by centrifugation in proteus clarification mini spin column (Protein Ark). The ssDNA containing filtrate was concentrated and buffer exchanged to annealing buffer (100 mM NaCl, 10 mM Tris pH 8.0, 1 mM EDTA), using amicon ultra 30 k centrifugal filters (Millipore). The template strand was made in a similar process, except that primers R3 and R2 were used in the first PCR step, and primer R5 was used in the second PCR step. The two ssDNA were annealed to form defective genes using the same annealing protocol. After annealing, the long defective genes were purified by Zeba spin desalting columns (ThermoFisher), eluting in TE buffer (10 mM Tris pH 8.0, 1 mM EDTA). They were further purified with Monarch PCR cleanup kit, eluting in nuclease free water.

### Fluorescence measurements

Epifluorescence microscopy (Leica DMI 8) was used to determine fluorescence intensity of reaction samples. For samples incubated in bulk solution, 200 nL of the solution was added to a lipid-in-oil solution in a PMMA chamber at the end of the incubation to determine the fluorescence intensity. The lipid was 1,2-diphytanoyl-sn-glycero-3-phosphocholine (DPhPC) or 1:1 (mass ratio) DPhPC:1-palmitoyl-2-oleoyl-glycero-3-phosphocholine (POPC), and the oil was 1:1 (v/v) hexadecane: silicone oil (AR20). Samples incubated in lipid-in-oil mixture or samples embedded in organogel were directly imaged under the microscope. The following excitation and emission filters were used. Alexa-488, DFHBI-1T, TO1B, mNeonGreen: excitation: 450-490 nm, emission: 500-550 nm; HBC620, mCherry: excitation 540-580 nm, emission 592-668 nm.

### RNA production from defective genes

To study the final fluorescence intensity when different concentrations of defective Broccoli and Broccoli signal were used, defective Broccoli (B1) and DNA signal (s1) were added to 10 mM Tris pH 8.0, 50 mM MgCl_2_, 1 mM DTT, 10 mM NTP (each), 60 μM DFHBI-1T, 50 mM KCl, and 5000 units mL^-1^ T7 RNA polymerase. The mixture was incubated at 37 °C for 3 h prior to fluorescence measurement. To study fluorescence activation of different fluorophores by different aptamers, 100 nM of either defective Broccoli (B1), defective Mango (M1), or defective Pepper (P2), were added to 100 nM DNA signal (s1), 20 mM MOPS pH 8.0, 50 mM MgCl_2_, 2 mM Spermidine, 1 mM DTT, 10 mM NTP (each), 50 mM KCl, and 5000 units mL^-1^ T7 RNA polymerase, in the presence of either 60 μM DFHBI-1T, 5 μM HBC620 or 2.5 μM TO1B. The mixture was incubated at 37 °C for 3 h prior to fluorescence measurement. To estimate strength of signal binding to defective genes, DNA melting temperature was calculated using online calculator from Promega, ^54^ with the following settings: 100 nM primer concentration, 10 mM MgCl_2_, and 50 mM KCl. To study selective activation of defective Broccoli with T7 promoter (B1) and defective Pepper with SP6 promoter (P1), 100 nM Broccoli signal (s2) and/or 100 nM Pepper signal (s3) were added to 5 mM MOPS pH 8.0, 50 mM MgCl_2_, 2 mM Spermidine, 1 mM DTT, 10 mM NTP (each), 50 mM KCl, 100 nM defective Broccoli (B1), 100 nM defective Pepper (P1), 60 μM DFHBI-1T, 5 μM HBC620, 5000 units mL^-1^ T7 RNA polymerase, and 2000 units mL^-1^ SP6 RNA polymerase. The mixture was incubated at 37 °C for 3 h prior to fluorescence measurement. To study selective activation of defective Mango with T7 promoter (M1) and defective Pepper with SP6 promoter (P1), 100 nM Mango signal (s4) and/or 100 nM Pepper signal (s3) were added to 20 mM MOPS pH 8.0, 50 mM MgCl_2_, 2 mM Spermidine, 1 mM DTT, 10 mM NTP (each), 50 mM KCl, 100 nM 11BM, 100 nM 31PPM, 60 μM DFHBI-1T, 5 μM HBC620, 2500 units mL^-1^ T7 RNA polymerase, and 2000 units mL^-1^ SP6 RNA polymerase. The mixture was incubated at 37 °C for 3 h prior to fluorescence measurement.

To study selective activation of defective Broccoli (B1) and defective Pepper (P2) with T7 promoter, 100 nM of either Broccoli signal (s2), Pepper signal (s5), or universal signal (s6) were added to 5 mM MOPS pH 8.0, 50 mM MgCl_2_, 1 mM DTT, 10 mM NTP (each), 50 mM KCl, 5000 units mL^-1^ T7 RNA polymerase, with either 100 nM defective Broccoli (B1), 1 uM Pepper signal inhibitor (I2) and 60 μM DFHBI-1T; or 100 nM defective Pepper (P2), 1 uM Broccoli signal inhibitor (I1) and 5 μM HBC620. The mixture was incubated at 37 °C for 3 h prior to fluorescence measurement. In the agarose gel electrophoresis analysis of strand displacement reaction, Pepper signal (s5) and defective Broccoli (B1) were first incubated at 37 °C for 30 minutes. Pepper signal inhibitor (I2) was then added and the mixture was incubated at 37 °C for a further 60 minutes before loading the samples onto agarose gel for analysis. To study selective activation of defective Mango (M1) and defective Pepper (P2) with T7 promoter, 100 nM of either Pepper signal (s5), Mango signal (s4), universal signal (s7), or polyA signal (s8), were added to 20 mM MOPS pH 8.0, 50 mM MgCl_2_, 1 mM DTT, 10 mM NTP (each), 50 mM KCl, 5000 units mL^-1^ T7 RNA polymerase, with either 100 nM defective Mango (M1), 1 uM Pepper signal inhibitor (I2) and 2.5 μM TO1B; or 100 nM defective Pepper (P2), 1 uM Mango signal inhibitor (M1) and 5 μM HBC620. The mixture was incubated at 37 °C for 3 h prior to fluorescence measurement. To study selective activation of defective Broccoli (B2), defective Mango (M1) and defective Pepper (P2) with T7 promoter, 100 nM of either Pepper signal (s5), or Mango signal (s4), or Broccoli signal (s9), were added to 20 mM MOPS pH 8.0, 50 mM MgCl_2_, 1 mM DTT, 10 mM NTP (each), 50 mM KCl, 5000 units mL^-1^ T7 RNA polymerase, with either 100 nM defective Mango (M1), 1 μM Pepper signal inhibitor (I2), 1 μM Broccoli signal inhibitor (I4), and 2.5 μM TO1B; or 100 nM defective Pepper (P2), 1 μM Mango signal inhibitor (M1), 1 μM Broccoli signal inhibitor (I4), and 5 μM HBC620; or 100 nM defective Broccoli (B2), 1 μM Mango signal inhibitor (M1), 1 μM Pepper signal inhibitor (I2), and 60 μM DFHBI-1T. The mixture was incubated at 37 °C for 3 h prior to fluorescence measurement.

To study fluorescence intensity of inverse defective Pepper (P3), 100 nM Pepper signal (s10) was mixed with 5 mM MOPS pH 8.0, 50 mM MgCl_2_, 2 mM spermidine, 1 mM DTT, 10 mM NTP (each), 50 mM KCl, 5000 units mL^-1^ T7 RNA polymerase, 5 μM HBC620, 100 nM inverse defective Pepper (P3), with or without 40,000 units mL^-1^ T4 DNA ligase. The mixture was incubated at 37 °C for 3 h prior to fluorescence measurement. To study fluorescence intensity of inverse defective Broccoli (B3), 100 nM Broccoli signal (s11) is mixed with 5 mM MOPS pH 8.0, 50 mM MgCl_2_, 2 mM spermidine, 1 mM DTT, 10 mM NTP (each), 50 mM KCl, 5000 units mL^-1^ T7 RNA polymerase, 60 μM DFHBI-1T, 100 nM inverse defective Broccoli (B3), with or without 40,000 units mL^-1^ T4 DNA ligase. The mixture was incubated at 37 °C for 3 h prior to fluorescence measurement. To study fluorescence intensity of logic gates, 1 μM each signal fragments were added to 5 mM MOPS pH 8.0, 50 mM MgCl_2_, 1 mM DTT, 10 mM NTP (each), 50 mM KCl, 5000 units mL^-1^ T7 RNA polymerase, 100 nM defective Broccoli (B4), and 60 μM DFHBI-1T, with or without 300,000 units mL^-1^ T3 DNA ligase. The mixture was incubated at 37 °C for 3 h prior to fluorescence measurement.

### Protein production from defective genes

To investigate the effect of signal concentration on the activation of defective mNG (G1), DNA signal (s12) and defective mNG (G1) were mixed with 1x PURExpress In Vitro Protein Synthesis Mix, and 1000 units mL^-1^ murine RNAse inhibitor. The mixture was incubated at 37 °C for 3 h prior to fluorescence measurement. To investigate DNA signal as a transcription factor mimic, 100 nM DNA signal (s12) was added to either 10 nM defective mCherry (C1), 10 nM defective mNG (G1), or 5 nM defective mCherry (C1) and 5 nM defective mNG (G1), with 1x PURExpress In Vitro Protein Synthesis Mix, and 1000 units mL^-1^ murine RNAse inhibitor. The mixture was incubated at 37 °C for 3 h prior to fluorescence measurement.

To activate αHL production using DNA signal in droplets, the recipient droplet (200 nL) contained 6 nM defective αHL (A1), 1x PURExpress In Vitro Protein Synthesis Mix, and 1000 units mL^-1^ murine RNAse inhibitor, with or without 20 nM DNA signal (s12). The reservoir droplet (200 nL) contained 1x solution A of PURExpress In Vitro Protein Synthesis Mix, 200 mM potassium monoglutamate, and 0.33 mM NBDG. The two droplets were separately pipetted into a lipid-in-oil solution with 10 mg/mL 1:1 (mass ratio) DPhPC: POPC in 1:1 (v/v) hexadecane: silicone oil (AR20). The droplets were gently brought together to allow lipid bilayer formation between the droplets. The constructs were then incubated at 37 °C and imaged periodically.

### Droplets with droplet-hydrogel bilayers

Droplets formed from droplet-hydrogel bilayers were constructed adapting protocol outlined in ^36^. The signal droplet (200 nL) contained 70 µM DNA signal (s6), 6.25 % glycerol, 1% ultra-low gelling agarose (melted), 5 mM MOPS pH 8.0, 50 mM MgCl_2_, 1 mM DTT, 10 mM NTP (each), 50 mM KCl, 2.5 μM TO1B and 0.1% F68. The receiver droplet (200 nL) contained 100 nM defective Mango (M1), 5000 units mL^-1^ T7 RNA polymerase, 1000 units mL^-1^ murine RNAse inhibitor, 1% ultra-low gelling agarose (melted), 5 mM MOPS pH 8.0, 50 mM MgCl_2_, 1 mM DTT, 10 mM NTP (each), 50 mM KCl, 2.5 μM TO1B and 0.1% F68. Similarly, the two cells were separately pipetted into a lipid-in-oil solution with 10 mg mL^-1^ 1:1 (mass ratio) DPhPC: POPC in 1:1 (v/v) hexadecane: silicone oil (AR20), gently brought together, and incubated for 10 minutes at 25 °C to form bilayer between the droplets. The droplets were then kept at 4 °C for 30 minutes, and activated by perfusing the lipid-in-oil solution with pure silicone oil (AR20) solution to break the bilayers. The constructs were kept at 4 °C for 60 minutes, and then incubated at 37 °C for 3 h prior to fluorescence imaging.

### Membrane potential mediated signaling in droplets

To characterize the translocation rate of ssDNA through different αHL mutant pores, the αHL monomers were expressed and purified with an established procedure. ^55^ Briefly, the mutants were constructed with Quikchange II site-directed mutagenesis kit (Agilent), following the manufacturer’s recommendations. The monomers were overexpressed in BL21(DE3)pLysS grown in LB media. The monomers were purified from the cell lysate by Ni-NTA column followed by size exclusion chromatography (HiLoad 26/600 Supedex 200 pg, GE Healthcare), eluting in 10 mM Tris pH 8.0, 150 mM NaCl. Translocation rate of ssDNA (s14) across different αHL mutants was studied using an established protocol. ^44^ Briefly, A planar bilayer of DPhPC was formed on a 100 μm diameter aperture in a 25 μm thick polytetrafluoroethylene film (Goodfellow Cambridge Limited) that divided a chamber into two compartments. Both compartments contained 1 M KCl, 25 mM Tris pH 8.0, and 100 μM EDTA. The protein and subsequently the ssDNA (s14) were added to the cis compartment, which was connected to ground. Planar bilayer current recordings were performed with a headstage (CV203BU, Axon Instruments), patch clamp amplifier (Axopatch 200B, Axon Instruments) and digitizer (Digidata 1550B, Molecular Devices).

To prepare droplets for DNA signal translocation across channel protein pores in droplet interface bilayers, 0.1 mm diameter Ag wires were oxidized in 15% sodium hypochlorite solution for at 25 °C for 15 minutes to form Ag/AgCl electrodes. The electrodes were coated twice with 1% Agarose in 100 mM KCl solution. The electrodes were attached onto male crimp (RS components), and connected to eONE amplifier (Elements) through a micromanipulator (NMN-21, Narishige). The tips of the electrodes were immersed in the lipid-in-oil solution of 10 mg mL^-1^ DPhPC in 1:1 (v/v) hexadecane: silicone oil (AR20). Signal droplets (200 nL) contained 100 mM NaCl, 0.2 mg mL^-1^ αHL monomer and 300 μM DNA signal (s13). Receiver droplets (200 nL) contained 100 mM NaCl. The two droplets were separately pipetted onto electrode tips in the lipid-in-oil solution, and gently brought together to form lipid bilayer between the droplets using the micromanipulator. The membrane potential was applied using Elements Data Reader software (Elements). The signaling droplet was held at ground, while the voltage in the receiver droplet alternated between +120 mV and −120 mV (2 s each). The bilayer conductance was measured using the seal test function, with the following parameters: Vpulse = 20 mV, Discard (% of Tpu) = 30.00, Tpulse = 1000 ms, Thold = 1000 ms. The number of proteins inserted into the bilayer was determined from the conductance. ^56^ Signaling was stopped when the number of proteins on the bilayer multiplied by the time elapsed reached 15,000 channel h (∼1.5 h). The two droplets were carefully separated, and the receiver droplet was supplemented to a final 1x PURExpress In Vitro Protein Synthesis mix, 1000 units mL^-1^ murine RNAse inhibitor and 3.33 ng µL^-1^ defective mCherry (C1). The receiver droplet was incubated at 37 °C for 3 h prior to fluorescence measurement.

## ASSOCIATED CONTENT

### Author Contributions

B.N., Y.Q. & H.B. designed and managed the project. B.N. performed all the experiments. B.N. and Y.D performed droplet electrophysiology experiments. B.N. and M.C. optimized the in vitro transcription protocols for aqueous-in-oil droplets. B.N. and R.K.K. performed the bilayer breaking experiments and analyzed all microscopy data. B.N. wrote the original draft of the manuscript. B.N., R.K.K., Y.D., Y.Q., H.B. revised the manuscript. All authors have given approval to the final version of the manuscript.

## ACKNOWLEDGMENT

This research was supported by a European Research Council Advanced Grant 833792 SYNTISU and a University of Oxford John Fell Fund (0007783). B.N. was supported by Croucher Scholarship. R.K.K. acknowledges funding from the Health Research Bridging Salary Scheme (0011044) and an EPSRC Open Plus Fellowship (EP/X010961/1). We acknowledge Michael Booth, Jorin Riexinger, Sejeong Lee, Juan Liu, and Denis Hartmann for their insightful discussions and contributions to other works related to this manuscript.

## ABBREVIATIONS

IVT: *in vitro* transcription
IVTT: in vitro transcription & translation

