## Supplementary for "Multifunctional Systems in Synthetic Biology: Single-Stranded DNA Signaling for Precise Control of Gene Activation"

Supplementary Text 1

Single-stranded region of defective genes

Most of the defective genes designed for this work (B1-2, B4, P1-2, M1, C1, G1, A1) have a single-stranded promoter, with the coding strand missing. As the signals bound to the defective genes, even if the signal complemented the defective gene perfectly, a nick was present between the 3' end of the signal and the defective gene. The strand break occurred downstream of the promoter on the coding strand, and therefore did not affect transcription. In fact, Martin group showed that full-length transcripts could still be made even when the coding strand was entirely missing after the promoter.^5^ We also designed defective genes with the template strand missing in the single-stranded promoter. Those defective gene produce low level of transcripts even in the presence of the complementary DNA signal, as DNA signal binding resulted in a nick (strand-break) on the template strand, which was less desirable. Nevertheless, we found that the use DNA ligase repaired the nick between the signal and inverse defective gene, improving the gene activation. (Figure S13, A to D).

**Supplementary Text 2**

Normalized fluorescence intensity

Small-molecule dyes exhibit varying levels of brightness when bound to their respective aptamers.^1^ To account for these differences, we normalized the fluorescence intensity in experiments involving the use of multiple dyes. The normalized fluorescence intensity of each sample was calculated using the following formula:

$\frac{I-I_{c}}{I_{max}-I_{c}}\times100$,

Where:

- *I* is the mean fluorescence intensity of the sample
- *I_c_* is the mean fluorescence intensity of the no-signal control
- *I_max_* is the mean fluorescence intensity of the brightest sample.

This normalization allows for easier comparison across different dyes, ensuring that variations in dye brightness do not influence the interpretation of experimental results. For further comparison, the fold-change in fluorescence intensity data is provided in the supplementary information.

**Supplementary Text 3**

Selective gene deactivation using T7 RNA polymerase only

Another strategy for selective gene activation involved using the same T7 RNA polymerase but with different adjacent sequences to the promoter for complementation of two different signals. It was more challenging to achieve signal specificity when only one polymerase was used, as all high efficiency T7 promoter variants share the same sequence from -17 to -5 of transcription start site (TAATACGACTCAC),^2^ where this conserved sequence has a calculated T_m_ of 47 ^o^C. Therefore, signal bound to the wrong defective gene would only detach above 47 ^o^C. At a transcription temperature of 37 ^o^C, all signals carrying the T7 promoter sequence would activate all defective genes controlled by the defective T7 promoter. To achieve selective gene activation using T7 RNA polymerase only, we constructed novel signals with toeholds that remained exposed after binding to the defective genes. Having exposed toeholds meant that the undesired signals could either bind directly to the complementary inhibitors or, upon binding to defective genes, be removed by the inhibitors through strand displacement reactions. (Figure S10, A and B).

Based on this principle, we designed a defective Broccoli (B1) and defective Pepper (P2) that had single-stranded T7 promoter sequences but were activated by different signals with toeholds as a consequence of different sequences flanking the promoter. Broccoli signal (s2) matched the single-stranded region on the defective Broccoli, with an extra 20 nucleotides (nt) at the 3' end of the signal. Similarly, Pepper signal (s5) matched the single-stranded region on defective Pepper, with an extra 20 nt at the 3' end. The Broccoli signal inhibitor (I1) and Pepper signal inhibitor (I2) were complementary to the Broccoli signal and the Pepper signal respectively. We also designed a universal signal (s6) which only had the complementary T7 promoter sequence, lacking the flanking sequences required for strand displacement reactions by the inhibitors.

In IVT mixtures that contained the defective Broccoli and the Pepper signal inhibitor, the Broccoli signal or the universal signal produced Broccoli aptamer, while the Pepper signal did not. Similarly, in IVT mixtures that contained the defective Pepper and the Broccoli signal inhibitor, the Pepper signal or the universal signal produced Pepper aptamer, while the Broccoli signal did not. In contrast, when inhibitors were not used, activation of the defective genes by the wrong signals were observed (Figure S10, C to H). To confirm that selective gene activation was facilitated by the strand displacement reaction, we verified that the Pepper signal could be removed from the defective Broccoli by the Pepper signal inhibitor, by analysing the size of the constructs with agarose gel electrophoresis (Figure S11). We showed the flexibility of this inhibitor strategy, which also worked for defective Mango and defective Pepper, with different strand displacement sequences compared to the defective Broccoli (Figure S12). In these experiments, the defective gene concentrations were 100 nM, the signal concentrations were 100 nM, and the signal inhibitor concentrations were 1 µM.

We then expanded the system to include three defective genes: defective Pepper (P2), defective Mango (M1), and defective Broccoli (B2). The Pepper signal (s5), Mango signal (s4), and Broccoli signal (s9) with toeholds were complementary to the Pepper signal inhibitor (I2), Mango signal inhibitor (I3), and Broccoli signal inhibitor (I4), respectively. We also designed a new universal signal (s7) where the sequences flanking the T7 promoter did not match any of the three inhibitors. In IVT mixtures of the defective Pepper (100 nM), the Mango signal inhibitor (1 µM) and the Broccoli signal inhibitor (1 µM), the defective Pepper was activated by the Pepper signal or the universal signal, but not by the Broccoli signal or the Mango signal. This was also observed when using the correct combinations of inhibitors for Broccoli and Mango signal/ defective gene pairs (Figure S13).

Willner^3^ and Winfree^4^ have also explored the use of inhibitor-mediated strand displacement to deactivate gene transcription. However, a key distinction between their system and ours lies in the effect on the promoter during gene regulation. Their approach involves the use of inhibitor strands to remove a segment of the 3'-5' strand of the promoter through a strand-displacement reaction. This method, while effective in halting gene activation, necessitates a strand break in the promoter, such as between positions -12 and -13.^3^ The requirement for a strand break within the promoter makes their system less robust, as such disruptions to the promoter can negatively impact RNA transcription and may not be permissible at all promoter positions. In contrast, our system offers a significant advantage by avoiding any strand breaks on the promoter when the signal binds to the defective gene. Instead, the strand break in our system occurs downstream of the promoter on the coding strand, and therefore does not affect transcription. In fact, Martin group showed that full-length transcripts could still be made even when the coding strand was entirely missing after the promoter.^5^ Since defects on the coding strand after the promoter are much more tolerable, our system is therefore a lot more robust to design, with the possibility of altering the length of the signal to shifting the location of the strand break to multiple locations.

Supplementary Text 4

ssDNA translocation rate across αHL pores

To improve the rate of ssDNA translocation across αHL pores, we constructed a new αHL mutant (E111N/N113R) to enhance the rate of DNA signal translocation in the presence of membrane potential. As αHL forms heptameric pores, the mutation removed 7 negative charges and introduced 7 positive charges into the pore lumen. At + 80 mV, the new mutant increased the rate of DNA translocation by 80 times compared to wild type αHL (wild type data reported^6^) (Figure S14).

Membrane potential across bilayers

In the signal translocation experiments across droplet-interface bilayers, The signalling droplet was held at ground (0 mV), while the voltage in the receiver droplet alternated between +120 mV and -120 mV every 2 seconds. This voltage protocol was chosen to avoid potential depletion of the Ag/AgCl electrodes and the droplet electrolytes.

T7 RNA polymerase activity affected by Ag/AgCl electrodes

The IVTT components were introduced after signal translocation across the bilayers. This was because T7 RNA polymerase within aqueous-in-oil droplet loses catalytic activity within 15 minutes of electrode insertion. The underlying cause of this phenomenon is under investigation.

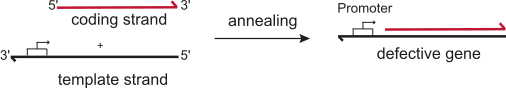

Figure S1. A schematic of the defective gene fabrication process. Defective genes were fabricated by annealing two single-stranded DNA of different lengths.

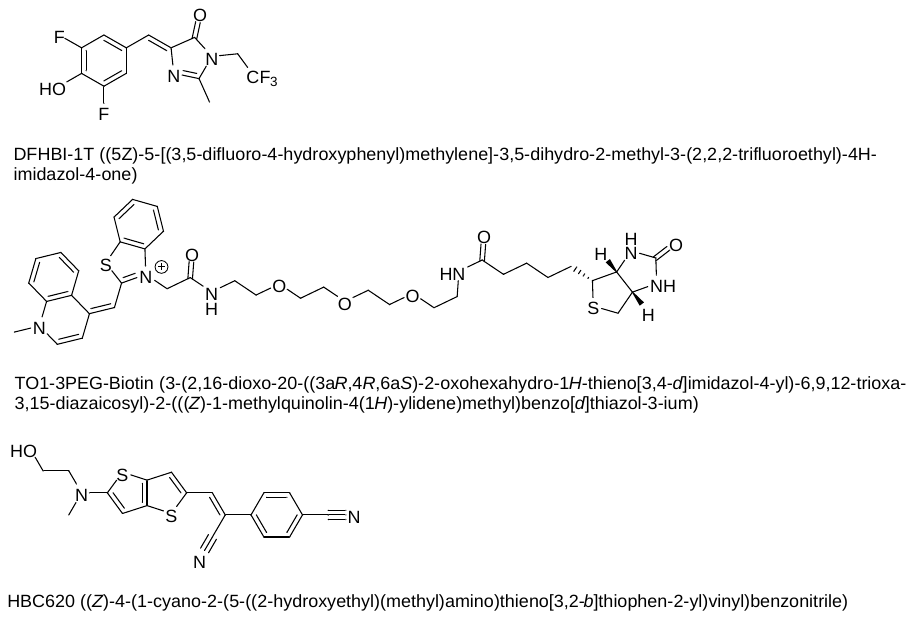

Figure S2. Structures and full chemical names of DFHBI-1T,^7^ TO1-3PEG-Biotin,^8^ and HBC620.^9^

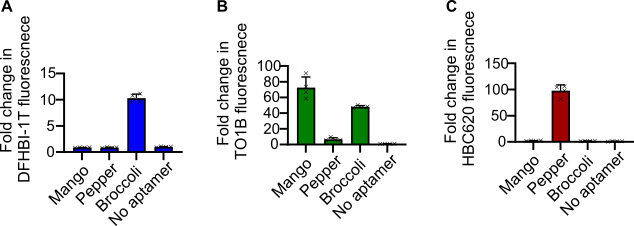

**Figure S3.** (**A to C**) Bar graphs of fold change in fluorescence intensities of (**A**) DFHBI-1T, (**B**) TO1B, and (**C**) HBC620 in an IVT mixture, where 100 nM DNA signal (s1) was added to 100 nM of either defective Broccoli (B1), defective Mango (M1), or defective Pepper (P2) after 3 h, compared to the control with no aptamer. Technical replicates are displayed by crosses and the heights of the bars are the mean. The error bars show the standard deviation.

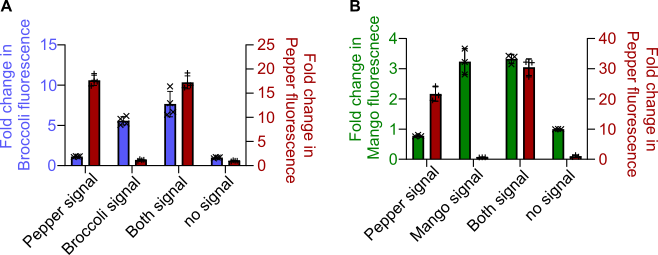

**Figure S4.** (**A**) A bar graph of fold change in fluorescence intensity of a defective Pepper (P1) and Broccoli (B1) mixture in an IVT mixture, 3 h after the addition of Pepper (s3) and/or Broccoli (s2) signals, compared to the control with no signal. The concentrations of signals and defective genes were 100 nM. (**B**) A bar graph of fold change in fluorescence intensity of a defective Mango (M1) and Pepper (P1) in an IVT mixture, 3 h after the addition of Pepper (s3) and/or Mango (s4) signals, compared to the control with no signal. In the bar graphs, technical replicates are displayed by crosses and the heights of the bars are the mean. The error bars show the standard deviation.

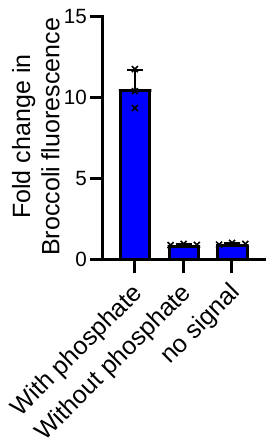

**Figure S5.** A bar graph of the fold change in fluorescence intensity of the Broccoli aptamer in an IVT mixture of the defective Broccoli, -10U signal fragment, and -10D signal fragment after 3 h, with or without 5’ phosphate on -10D, compared to the control with no signal. -10U and -10D were the upstream and downstream signal fragments from splitting the Broccoli signal after position -10. Technical replicates are displayed by crosses and the heights of the bars are the mean. The error bars show the standard deviation.

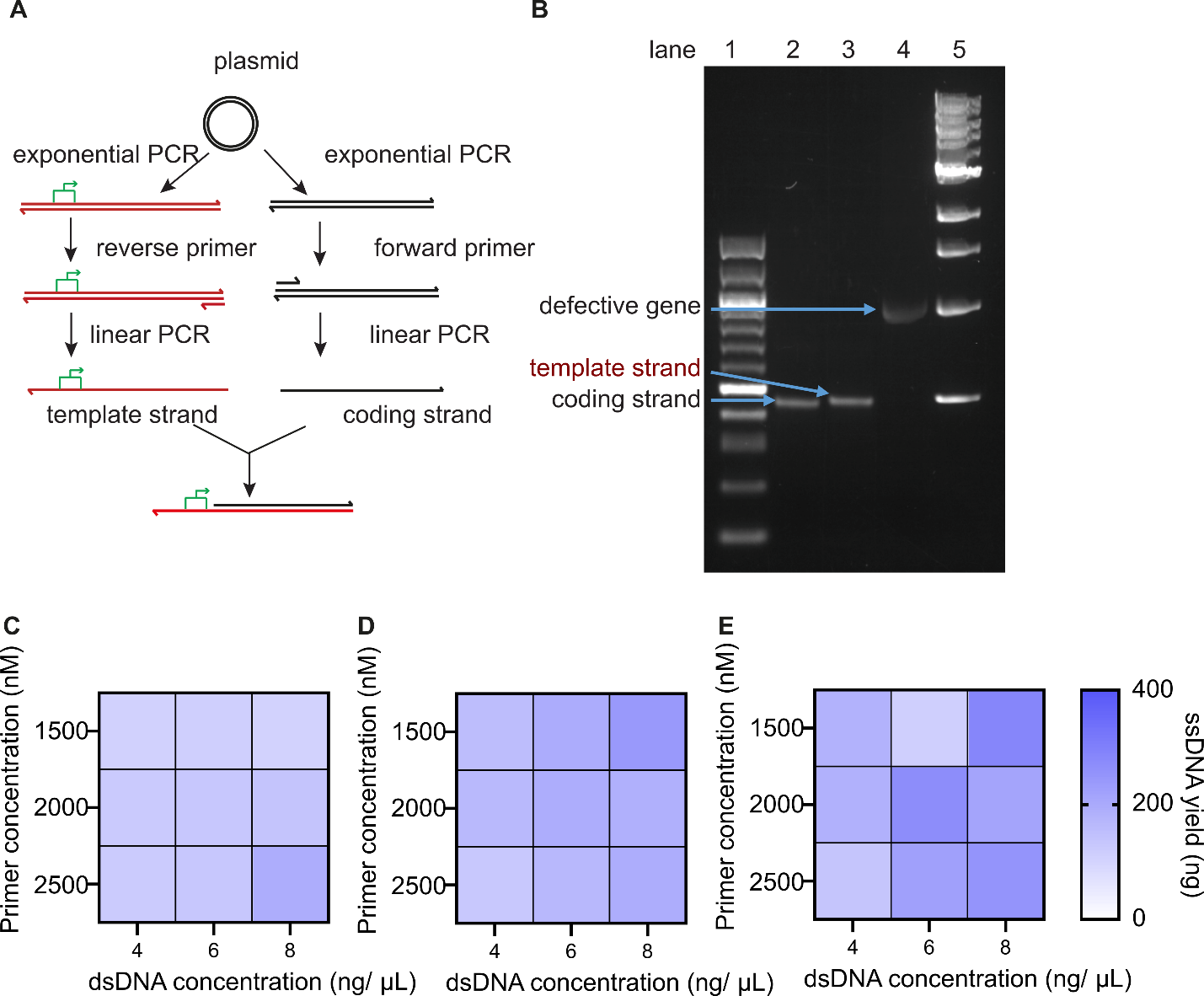

**Figure S6.** Long defective gene (>1000 bp) synthesis. (A) A schematic representation. Long ssDNA (>1000 nt) was made by 2-step PCR. The required sequences was first amplified by standard exponential PCR with two primers. The exponential PCR product was then used to produce ssDNA in a linear PCR reaction, where only one primer was present. Defective genes encoding protein sequences (>1000 bp) were made by annealing two ssDNA of different lengths. (B) Agarose gel (1%) electrophoresis of DNA constructs. Lane 1: DNA ladder 100 bp (NEB). Lane 2: Coding strand of defective mNG. Lane3: Template strand of defective mNG. Lane 4: Defective mng formed from annealing the two ssDNA. Lane 5: DNA ladder 1 kb (NEB). (C to E) Optimization of linear PCR reaction to produce ssDNA used in the defective gene synthesis. A heat map of mean ssDNA yield (ng) of coding strand of defective mNG per 40 µL PCR reaction mixture at different primer concentrations, dsDNA concentrations and number of thermocycles. (C) 10 cycles, (D) 15 cycles, (E) 20 cycles. Values in the heatmap are the mean of n = 2 technical repeats.

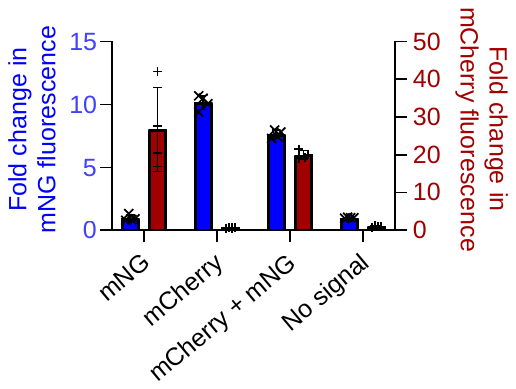

**Figure S7.** A bar graph of fold change in fluorescence from an IVTT mixture containing the DNA signal (s12) and different defective genes 3 h after signal addition, compared to the no signal control. In the bar graphs, technical replicates are displayed by crosses and the heights of the bars are the mean. The error bars show the standard deviation.

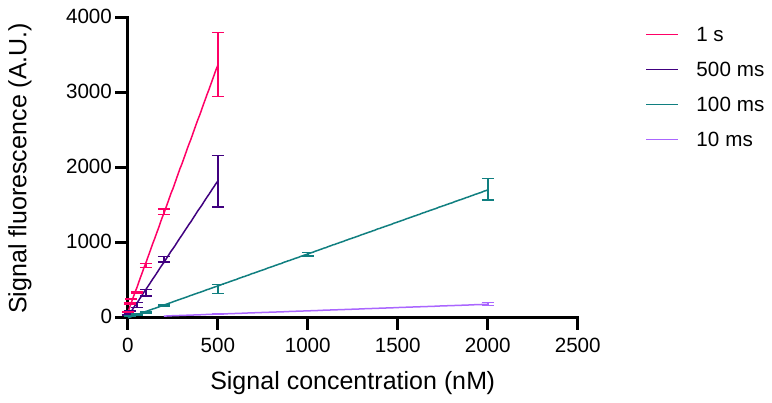

**Figure S8.** Calibration curve relating measured fluorescence intensity of Alexa-488 labelled signal to signal concentration. The best fit line was constructed using the means of fluorescence measurements, and the error bars represent standard deviation (n = 3).

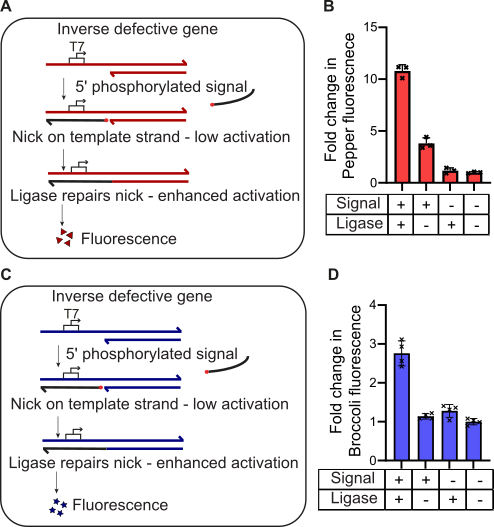

**Figure S9.** Inverse defective gene activation with DNA signal and DNA ligase. (**A**) A schematic of inverse defective Pepper activation. Inverse defective Pepper with SP6 promoter produced low level of transcripts even in the presence of DNA signal, as DNA signal binding left a nick (strand-break) on the template strand in the SP6 promoter. DNA ligase repaired the nick between the signal and inverse defective gene and improved gene activation. (**B**) A bar graph of fold change in fluorescence intensity of the Pepper aptamer in a bulk mixture of Pepper signal (s10) and inverse defective Pepper (P3) after 3 h, with or without DNA ligase, compared to the control with no signal. (**C**) A schematic of inverse Broccoli activation. Inverse defective Broccoli with T7 promoter produced low level of transcripts even in the presence of DNA signal, as DNA signal binding leaves a nick (strand-break) on the template strand in the T7 promoter region. (D) A bar graph of fold change in fluorescence intensity of the Broccoli aptamer in a bulk mixture of Broccoli signal (s11) and inverse defective Broccoli (B3) after 3 h, with or without T4 DNA ligase, compared to the control with no DNA signal. In the bar graphs, technical replicates are displayed by crosses and the heights of the bars are the mean. The error bars show the standard deviation.

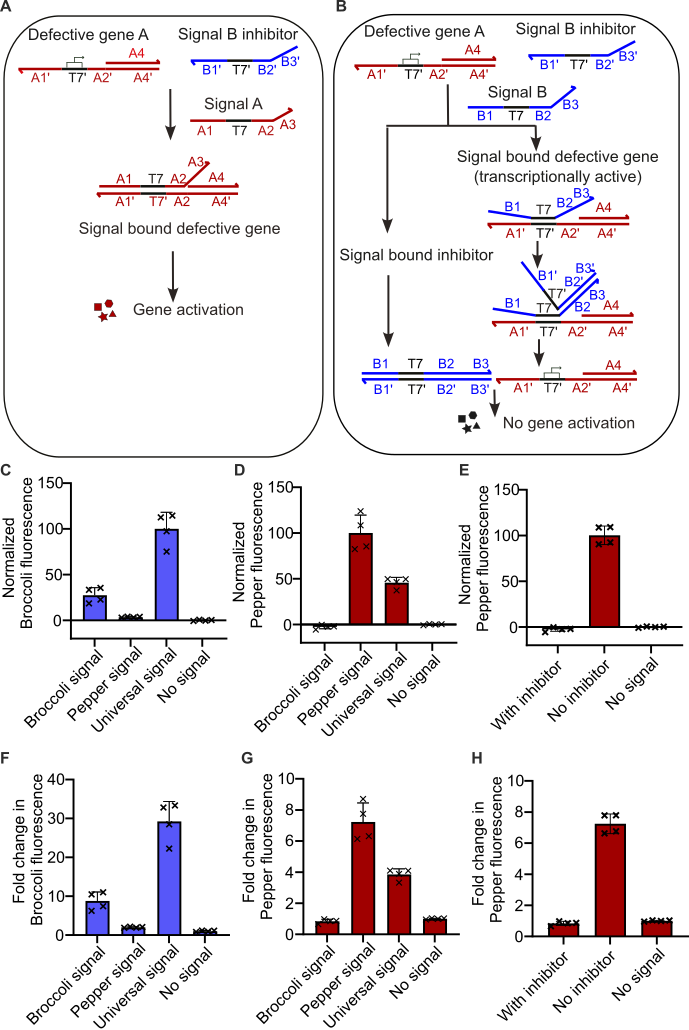

**Figure S10.** DNA signals selectively activate genes transcribed by the same polymerase. **(A)** A schematic of successful gene activation using an uninhibited signal. The 5'-3' strand and 3'-5' strand sequences of the T7 promoter are labelled T7 and T7' respectively. Other than the promoter sequences, DNA sequences on the coding strand of defective gene A are labelled A1, A2, A3 etc. DNA sequences on the template strand of defective gene A are numbered A1', A2', A3' etc. Sequences with labels differ by ' (e.g. A1 and A1') are reverse complementary to each other. In a mixture of defective gene A and signal B inhibitor, signal A is not inhibited by signal B inhibitor and can activate defective gene A. Signal A matches the single-stranded region on defective gene A, with an extra 20 nucleotides (nt) on the 3' end of the signal. As signal A is longer than the single-stranded region on defective gene A, a toehold (A3) is exposed after signal binding. Defective gene A is activated as the presence of the toehold does not inhibit transcription. (**B**) Schematic of signal removal using signal inhibitors. In a mixture of defective gene A and signal B inhibitor, signal B can bind directly to signal B inhibitor as they are complementary to each other, or it can bind to defective gene A due to the T7 promoter sequence. In the latter case, the exposed toehold (B3) is recognized by signal B inhibitor, and signal B inhibitor removes signal B from defective gene A. In either case, defective gene A is not activated by signal B. (**C**) A bar graph of normalized fluorescence intensity of the Broccoli aptamer in an IVT mixture of the defective Broccoli (B1) and the Pepper signal inhibitor (I2) 3 h after Broccoli (s2), Pepper (s5), or Universal (s6) signal addition. The Universal signal did not have flanking sequences to the promoter and was not removed by the inhibitors. The mean fluorescence intensity of the no-signal control and the brightest sample were normalized to 0 and 100 respectively. (**D**) A bar graph of normalized fluorescence intensity of the Pepper aptamer in an IVT mix of the defective Pepper gene (P2) and Broccoli signal inhibitor (I1) 3 h after signal addition. The mean fluorescence intensity of the no-signal control and the brightest sample were normalized to 0 and 100 respectively. (**E**) A bar graph of normalized fluorescence intensity of the Pepper aptamer in an IVT mixture of the defective Pepper (P2) and the Broccoli signal (s2) after 3 h, with or without Broccoli signal inhibitor (I1). The mean fluorescence intensity of the no signal control and the brightest sample were normalized to 0 and 100 respectively. (**F**) A bar graph of fold change in fluorescence intensity of the Broccoli aptamer in an IVT mixture of the defective Broccoli (B1) and the Pepper signal inhibitor (I2), 3 h after Broccoli (s2), Pepper (s5), or Universal (s6) signal addition, compared to the control with no signal. The Universal signal did not have flanking sequences to the promoter and was not removed by the inhibitors. (**G**) A bar graph of fold change in fluorescence intensity of the Pepper aptamer in an IVT mixture of the defective Pepper gene (P2) and Broccoli signal inhibitor (I1) 3 h after signal addition, compared to the control with no signal. (**H**) A bar graph of normalized fluorescence intensity of the Pepper aptamer in an IVT mixture of the defective Pepper (P2) and the Broccoli signal (s2) after 3 h, with or without Broccoli signal inhibitor (I1), compared to the control with no signal. In the bar graphs, technical replicates are displayed by crosses and the heights of the bars are the mean. The error bars show the standard deviation. The defective gene concentrations were 100 nM, the signal concentrations were 100 nM, and the signal inhibitor concentrations were 1 µM.

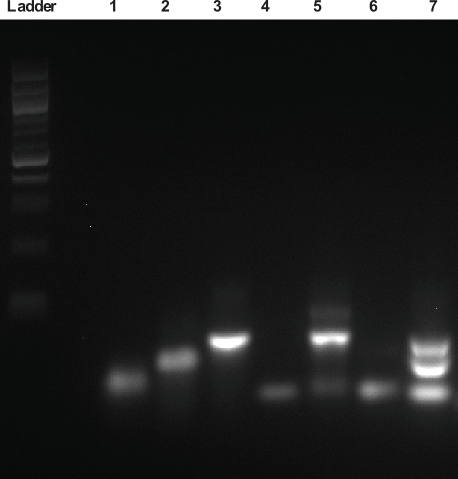

**Figure S11.** Agarose gel electrophoresis of DNA constructs (3% Agarose). Ladder: 100 bp DNA ladder (NEB). Lane 1: Coding strand of defective Broccoli (B1) (49 nt). Lane 2: template strand of defective Broccoli (B1) (74 nt). Lane 3: Defective broccoli (B1, 49 bp and 25 nt overhang), formed by annealing the coding strand and template strand. Lane 4: Pepper Signal (s5, 44 nt). Lane 5: Pepper signal (s5, 44 nt) bound to defective Broccoli (B1, 49 bp and 25 nt overhang). Lane 6: Pepper signal inhibitor (I2, 44 nt). Lane 7: Pepper signal inhibitor (I2) removes Pepper signal (s5) from defective Broccoli (B1). nt: nucleotides, bp: base pairs.

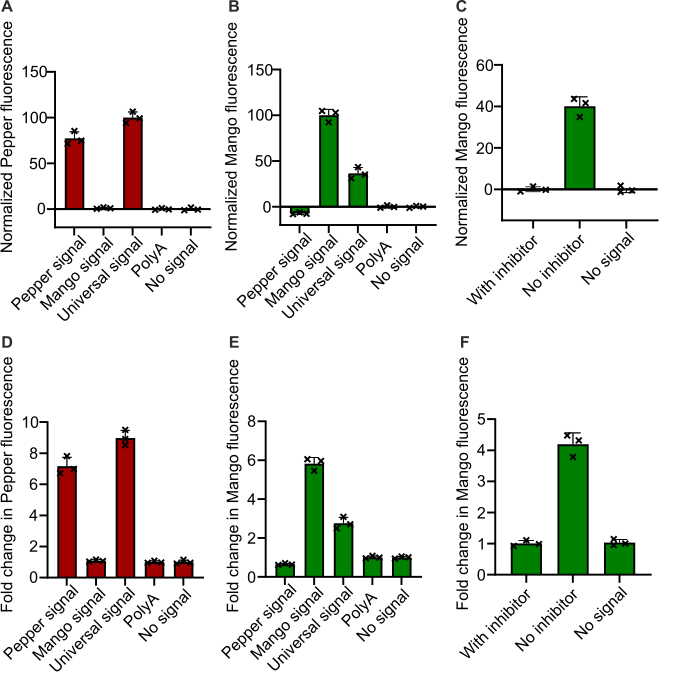

**Figure S12.** (**A**) A bar graph of normalized fluorescence intensity of the Pepper aptamer in an IVT mixture of defective Pepper (P2) and Mango signal inhibitor (I3) 3 h after signal addition. The signals were Pepper signal (s5), Mango signal (s4), Universal signal (s7) or PolyA control (s8). (**B**) A bar graph of normalized fluorescence intensity of the Mango aptamer in an IVT mixture of the defective Mango (M1) and Pepper signal inhibitor (I2) 3 h after signal addition. The mean fluorescence intensity of the no-signal control and the brightest sample were normalized to 0 and 100 respectively. (**C**) A bar graph of fluorescence intensity of the Mango aptamer in an IVT mixture of defective Mango (M1) and Pepper signal (s5) after 3 h, with or without Pepper signal inhibitor (I2). For each aptamer, the mean fluorescence intensity of the no signal control and the brightest sample were normalized to 0 and 100 respectively. (**D**) A bar graph of fluorescence intensity of the Pepper aptamer in an IVT mixture of defective Pepper (P2) and Mango signal inhibitor (I3) after 3 h signal addition, compared to the control with no signal. (**E**) A bar graph of the fold change in fluorescence intensity of the Mango aptamer in an IVT mixture of the defective Mango (M1) and Pepper signal inhibitor (I2) after 3 h of signal addition, compared to the control with no signal. (**F**) A bar graph of fluorescence intensity of the Mango aptamer in a bulk mixture of defective Mango (M1) and Pepper signal (s5) after 3 h, with or without Pepper signal inhibitor (I2), compared to the control with no signal. In the bar graphs, technical replicates are displayed by crosses and the heights of the bars are the mean. The error bars show the standard deviation. The defective gene concentrations were 100 nM, the signal concentrations were 100 nM, and the signal inhibitor concentrations were 1 µM.

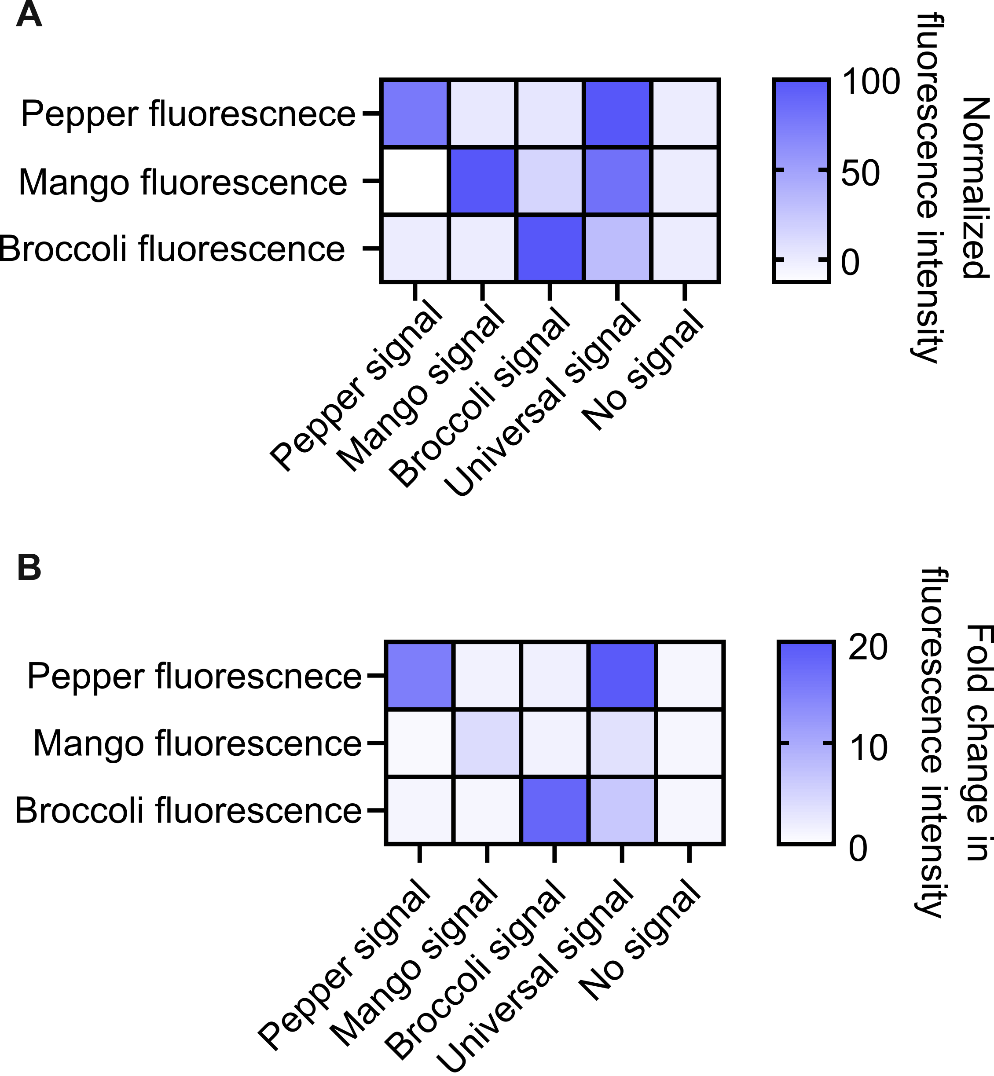

**Figure S13.** (**A**) A heat map of normalized fluorescence intensity of RNA aptamers 3 h after various signals were added. The first row shows normalized fluorescence intensity of the Pepper aptamer in an IVT mixture of the defective Pepper (P2), the Mango signal inhibitor (I3), and the Broccoli signal inhibitor (I4). The second and third rows show normalized fluorescence intensity of the Mango aptamer and the Broccoli aptamer respectively, in an IVT mixture of the correct combination of defective genes and signal inhibitors. For each aptamer, the mean fluorescence intensity of the no-signal control and the brightest sample were normalized to 0 and 100 respectively. In the bar graphs, technical replicates are displayed by crosses and the heights of the bars are the mean. The error bars show the standard deviation, (**B**) A heat map of fold change in fluorescence intensity of RNA aptamers 3 h after various signals were added, compared to the control with no signal. The first row shows fluorescence intensity of the Pepper aptamer in an IVT mixture of the defective Pepper (P2), the Mango signal inhibitor (I3), and the Broccoli signal inhibitor (I4). The second and third rows show fluorescence intensity of the Mango aptamer and the Broccoli aptamer respectively, in an IVT mixture of the correct combination of defective genes and signal inhibitors. The defective gene concentrations were 100 nM, the signal concentrations were 100 nM, and the signal inhibitor concentrations were 1 µM.

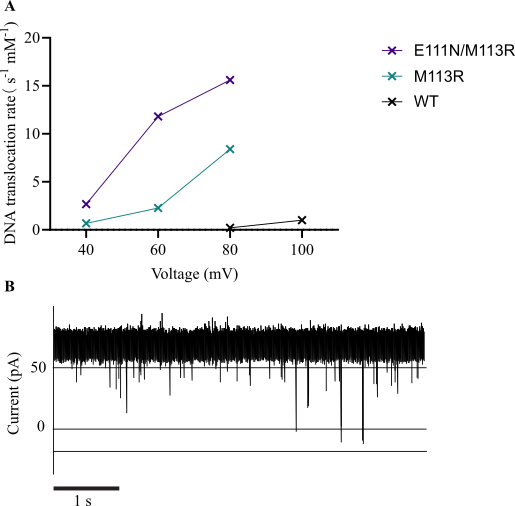

**Fig S14.** (**A**) DNA translocation rate (s14) across different αHL pores. The crosses are the mean translocation rates from more than 30 translocation events, the error bars are the standard error of the mean. Data for wild type αHL was reported^47^. (**B**) Representative current trace of DNA translocation through single E111N/M113R pore at +60 mV.

| Name | Sequence |
| --- | --- |
| s1 | Gaatttaatacgactcac |
| s2 | Gaatttaatacgactcacaatcgggagacggtcgggtccagata |
| s3 | Gaattatttaggtgacactatagaaatccccaatcgtggcgtg |
| s4 | Gaatttaatacgactcacaatcggggtacgaaggaagg |
| s5 | Cagtctaatacgactcactccgggatccccaatcgtggcgtgtc |
| s6 | Taatacgactcac |
| s7 | Atccgtaatacgactcaccgagggagacggtcgggtccagata |
| s8 | Aaaaaaaaaaaaaaaaaaaaaaaaaaaaaaaaaaaaaaaa |
| s9 | Tctgataatacgactcactatagggagacggtcgggtccag |
| s10 | [phos] gccacgattggggatttctatagtgtcacctaaataattc |
| s11 | [phos] gacccgaccgtctcccgattgtgagtcgtattaaattc |
| s12 | Gaaattaatacgactcactatagggtctagaaataatttt |
| s13 | [Alexa-488] gaaattaatacgactcactatagggtctagaaataatttt |
| s14 | Aaaaaaaaaaaaaaaaaaaaattcccccccccccccccccccccttaaaaaaaaaattccccccccccttaaaaaaaaaattcccccccccc |
| -12U | Gaatttaatac |
| -12D | [phos] gaactcacaatcgggagacggtcgggtccagata |
| -11U | Gaatttaatacg |
| -11D | [phos] actcacaatcgggagacggtcgggtccagata |
| -10U | Gaatttaatacga |
| -10D | [phos] ctcacaatcgggagacggtcgggtccagata |
| -9U | Gaatttaatacgac |
| -9D | [phos] tcacaatcgggagacggtcgggtccagata |
| I1 | Tatctggacccgaccgtctcccgattgtgagtcgtattaaattc |
| I2 | Gacacgccacgattggggatcccggagtgagtcgtattagactg |
| I3 | Ccttccttcgtaccccgattgtgagtcgtattaaattc |
| I4 | ctggacccgaccgtctcCCTATAGTGAGTCGTATTATCAGA |
| B1 | gaatttaatacgactcacaatcgggAGACGGTCGGGTCCAGATATTCGTATCTGTCGAGTAGAGTGTGGGCTCC |
| B2 | tctgataatacgactcactatagggAGACGGTCGGGTCCAGATATTCGTATCTGTCGAGTAGAGTGTGGGCTCC |
| B3 | gaatttaatacgactcacaatcgggagacggtcgggtcCAGATATTCGTATCTGTCGAGTAGAGTGTGGGCTCC |
| B4 | gaatttaatacgactcacaatcgggagacggtcgggtccagataTTCGTATCTGTCGAGTAGAGTGTGGGCTCC |
| P1 | gaattatttaggtgacactatagaaatccccaatcgtggcgtgTCGGCCTGCTTCGGCAGGCACTGGCGCCGGGATTTC |
| P2 | cagtctaatacgactcactccgggATCCCCAATCGTGGCGTGTCGGCCTCTCCCAATCGTGGCGTGTCGGCCTCTCTTCGGAGAGGCACTGGCGCCGGAGAGGCACTGGCGCCGGGATC |
| P3 | gaattatttaggtgacactatagaaatccccaatcgtggcGTGTCGGCCTGCTTCGGCAGGCACTGGCGCCGGGATTTC |
| M1 | gaatttaatacgactcacaatcggggTACGAAGGAAGGTTTGGTATGTGGTATATTCGTAC |
| R1 | gtttaactttaagaaggaggtatacatatggtgag |
| R2 | gatatagttcctcctttcagcaaaaaacccctca |
| R3 | gaaattaatacgactcactatagggtctagaaa |
| R4 | gtttaactttaagaaggaggtatac |
| R5 | gatatagttcctcctttcagcaaaaaacc |
| C1 | gaaattaatacgactcactatagggtctagaaataattttGTTTAACTTTAAGAAGGAGGTATACATATGGTGAGCAAGGGCGAGGAGGATAACATGGCCATCATCAAGGAGTTCATGCGCTTCAAGGTGCACATGGAGGGCTCCGTGAACGGCCACGAGTTCGAGATCGAGGGCGAGGGCGAGGGCCGCCCCTACGAGGGCACCCAGACCGCCAAGCTGAAGGTGACCAAGGGTGGCCCCCTGCCCTTCGCCTGGGACATCCTGTCCCCTCAGTTCATGTACGGCTCCAAGGCCTACGTGAAGCACCCCGCCGACATCCCCGACTACTTGAAGCTGTCCTTCCCCGAGGGCTTCAAGTGGGAGCGCGTGATGAACTTCGAGGACGGCGGCGTGGTGACCGTGACCCAGGACTCCTCCCTGCAGGACGGCGAGTTCATCTACAAGGTGAAGCTGCGCGGCACCAACTTCCCCTCCGACGGCCCCGTAATGCAGAAGAAGACCATGGGCTGGGAGGCCTCCTCCGAGCGGATGTACCCCGAGGACGGCGCCCTGAAGGGCGAGATCAAGCAGAGGCTGAAGCTGAAGGACGGCGGCCACTACGACGCTGAGGTCAAGACCACCTACAAGGCCAAGAAGCCCGTGCAGCTGCCCGGCGCCTACAACGTCAACATCAAGTTGGACATCACCTCCCACAACGAGGACTACACCATCGTGGAACAGTACGAACGCGCCGAGGGCCGCCACTCCACCGGCGGCATGGACGAGCTGTACAAGTAATGAGGATCCCGGGAATTCTCGAGTAAGGTTAACCTGCAGGAGGCCTTTAATTAAGGTGGTGCGGCCGCGCTAGCGGTCCCGGGGGATCGATCCGGCTGCTAACAAAGCCCGAAAGGAAGCTGAGTTGGCTGCTGCCACCGCTGAGCAATAACTAGCATAACCCCTTGGGGCCTCTAAACGGGTCTTGAGGGGTTTTTTGCTGAAAGGAGGAACTATATC |
| G1 | gaaattaatacgactcactatagggtctagaaataattttGTTTAACTTTAAGAAGGAGGTATACATATGGTGAGCAAGGGCGAGGAGGATAACATGGCCTCTCTCCCAGCGACACATGAGTTACACATCTTTGGCTCCATCAACGGTGTGGACTTTGACATGGTGGGTCAGGGCACCGGCAATCCAAATGATGGTTATGAGGAGTTAAACCTGAAGTCCACCAAGGGTGACCTCCAGTTCTCCCCCTGGATTCTGGTCCCTCATATCGGGTATGGCTTCCATCAGTACCTGCCCTACCCTGACGGGATGTCGCCTTTCCAGGCCGCCATGGTAGATGGCTCCGGATACCAAGTCCATCGCACAATGCAGTTTGAAGATGGTGCCTCCCTTACTGTTAACTACCGCTACACCTACGAGGGAAGCCACATCAAAGGAGAGGCCCAGGTGAAGGGGACTGGTTTCCCTGCTGACGGTCCTGTGATGACCAACTCGCTGACCGCTGCGGACTGGTGCAGGTCGAAGAAGACTTACCCCAACGACAAAACCATCATCAGTACCTTTAAGTGGAGTTACACCACTGGAAATGGCAAGCGCTACCGGAGCACTGCGCGGACCACCTACACCTTTGCCAAGCCAATGGCGGCTAACTATCTGAAGAACCAGCCGATGTACGTGTTCCGTAAGACGGAGCTCAAGCACTCCAAGACCGAGCTCAACTTCAAGGAGTGGCAAAAGGCCTTTACCGATGTGATGGGCATGGACGAGCTGTACAAGTAATGAGGATCCCGGGAATTCTCGAGTAAGGTTAACCTGCAGGAGGCCTTTAATTAAGGTGGTGCGGCCGCGCTAGCGGTCCCGGGGGATCGATCCGGCTGCTAACAAAGCCCGAAAGGAAGCTGAGTTGGCTGCTGCCACCGCTGAGCAATAACTAGCATAACCCCTTGGGGCCTCTAAACGGGTCTTGAGGGGTTTTTTGCTGAAAGGAGGAACTATATC |
| A1 | gaaattaatacgactcactatagggtctagaaataattttGTTTAACTTTAAGAAGGAGGTATACATATGGCAGATTCTGATATTAATATTAAAACCGGTACTACAGATATTGGAAGCAATACTACAGTAAAAACAGGTGATTTAGTCACTTATGATAAAGAAAATGGCATGCACAAAAAAGTATTTTATAGTTTTATCGATGATAAAAATCACAATAAAAAACTGCTAGTTATTAGAACAAAAGGTACCATTGCTGGTCAATATAGAGTTTATAGCGAAGAAGGTGCTAACAAAAGTGGTTTAGCCTGGCCTTCAGCCTTTAAGGTACAGTTGCAACTACCTGATAATGAAGTAGCTCAAATATCTGATTACTATCCAAGAAATTCGATTGATACAAAAAACTATATGAGTACTTTAACTTATGGATTCAACGGTAATGTTACTGGTGATGATACAGGAAAAATTGGCGGCCTTATTGGTGCAAATGTTTCGATTGGTCATACACTGAACTATGTTCAACCTGATTTCAAAACAATTTTAGAGAGCCCAACTGATAAAAAAGTAGGCTGGAAAGTGATATTTAACAATATGGTGAATCAAAATTGGGGACCATACGATCGAGATTCTTGGAACCCGGTATATGGCAATCAACTTTTCATGAAAACTAGAAATGGTTCTATGAAAGCAGCAGATAACTTCCTTGATCCTAACAAAGCAAGTTCTCTATTATCTTCAGGGTTTTCACCAGACTTCGCTACAGTTATTACTATGGATAGAAAAGCATCCAAACAACAAACAAATATAGATGTAATATACGAACGAGTTCGTGATGATTACCAATTGCATTGGACTTCAACAAATTGGAAAGGTACCAATACTAAAGATAAATGGACAGATCGTTCTTCAGAAAGATATAAAATCGATTGGGAAAAAGAAGAAATGACAAATTAATGAGGATCCCGGGAATTCTCGAGTAAGGTTAACCTGCAGGAGGCCTTTAATTAAGGTGGTGCGGCCGCGCTAGCGGTCCCGGGGGATCGATCCGGCTGCTAACAAAGCCCGAAAGGAAGCTGAGTTGGCTGCTGCCACCGCTGAGCAATAACTAGCATAACCCCTTGGGGCCTCTAAACGGGTCTTGGAGGGGTTTTTTGCTGAAAGGAGGAACTATATC |

**Table S1.** DNA sequences. All sequences are from 5' to 3'. Lower case denotes single-stranded sequences, upper case denotes double-stranded sequences. [phos]: phosphorylated 5' end. [Alexa-488]: Alexa-488 modified 5' end.
